# The neural dynamics of hierarchical Bayesian inference in multisensory perception

**DOI:** 10.1101/504845

**Authors:** Tim Rohe, Ann-Christine Ehlis, Uta Noppeney

## Abstract

Transforming the barrage of sensory signals into a coherent multisensory percept relies on solving the binding problem – deciding whether signals come from a common cause and should be integrated, or instead be segregated. Human observers typically arbitrate between integration and segregation consistent with Bayesian Causal Inference, but the neural mechanisms remain poorly understood. We presented observers with audiovisual sequences that varied in the number of flashes and beeps. Combining Bayesian modelling and EEG representational similarity analyses, we show that the brain initially represents the number of flashes and beeps and their numeric disparity mainly independently. Later, it computes them by averaging the forced-fusion and segregation estimates weighted by the probabilities of common and independent cause models (i.e. model averaging). Crucially, prestimulus oscillatory alpha power and phase correlate with observers’ prior beliefs about the world’s causal structure that guide their arbitration between sensory integration and segregation.

---

In everyday life, the brain is constantly confronted with a myriad of sensory signals. Imagine you are skipping stones on a lake. Each time the stone bounces off the water’s surface, you see the impact and hear a brief splash. Should you integrate or segregate signals from vision and audition to estimate how many times the stone hits the water’s surface? Hierarchical Bayesian Causal Inference provides a rational strategy to arbitrate between information integration and segregation by explicitly modelling the underlying potential causal structures, i.e. whether visual impacts and splash sounds are caused by common or independent events^1,2^. Under the assumption of a common cause, signals are integrated weighted by their relative precisions (or reliabilities, i.e. the reciprocal of variance) into one single ‘forced-fusion’ numeric estimate^3,4^. If, however, some splash sounds are caused by a stone hitting the water surface out of the observer’s sight (e.g. another person throwing a stone), audition and vision will provide conflicting information. In this segregation case, the brain needs to estimate the number of events independently for vision and audition. Importantly, the brain cannot directly access the world’s causal structure, but needs to infer it from the signals’ noisy sensory representations based on correspondence cues such as temporal synchrony or spatial co-location. To account for observers’ causal uncertainty, a final Bayesian Causal Inference estimate is computed by combining the ‘forced-fusion’ and the task-relevant unisensory segregation estimates weighted by the posterior probability of common or independent causes^1^. Perception thus relies crucially on inferring the hidden causal structure that generated the sensory signals.

Accumulating evidence suggests that human and animal observers arbitrate between sensory integration and segregation approximately in line with Bayesian Causal Inference^1,5-8^. For small intersensory conflicts, when it is likely that signals come from a common cause, observers integrate sensory signals approximately weighted by their relative precisions^3,4,9^, which leads to intersensory biases and perceptual illusions. Most prominently, in the sound-induced flash illusion, observers tend to perceive two flashes when a single flash appears together with two sounds^10^. For large intersensory conflicts such as temporal asynchrony, spatial disparity or numeric disparity, multisensory integration breaks down and crossmodal biases are attenuated^5,11^.

At the neural level, a recent fMRI study has demonstrated that the human brain performs multisensory Bayesian Causal Inference for spatial localization by encoding multiple spatial estimates across the cortical hierarchy^12,13^. While low-level sensory areas represented spatial estimates mainly under the assumption of separate causes, posterior parietal areas integrated sensory signals under the assumption of a common cause. Only at the top of the cortical hierarchy, in anterior parietal areas, the brain formed a Bayesian Causal Inference estimate that takes into account the observers’ uncertainty about the signals’ causal structure.

In summary, the brain should entertain two models of the sensory inputs, namely that the inputs are generated by common (i.e. forced-fusion model) or independent sources (i.e. segregation model). Using a decisional strategy called model averaging, hierarchical Bayesian Causal Inference accounts for the brain’s uncertainty about the world’s causal structure by averaging the forced-fusion and the task-relevant unisensory segregation estimates weighted by the posterior probabilities of their respective causal structures. Hence, hierarchical Bayesian Causal Inference goes beyond estimating an environmental property (e.g. numerosity, location) and involves inferring a causal model of the world (i.e. structure inference).

The hierarchical nature of Bayesian Causal Inference raises the intriguing question of how these computations evolve dynamically over time in the human brain. To assess this, we fitted the Bayesian Causal Inference model to observers’ behavioral responses and then investigated how observers’ forced-fusion, the full-segregation auditory and visual estimates and the final Bayesian Causal Inference (i.e. model averaging) estimates are dynamically encoded in neural responses measured with EEG. While the brain is likely to update all estimates continuously in recurrent loops across the cortical hierarchy^14,15^, the neural representations of unisensory segregation and forced-fusion estimates may be more pronounced at earlier latencies than the final Bayesian Causal Inference estimate whose computation requires the posterior probabilities of the potential causal structures (i.e. common vs. independent causes). Moreover, neural activity (i.e. alpha-, beta-and gamma-oscillations^16,17^) prior to stimulus onset may modulate the causal prior or precision of sensory representations (e.g. visual variance) in early visual cortices and thereby in turn influence the outcome of Bayesian Causal Inference. We combined psychophysics, computational modelling and EEG representational similarity analyses to characterize the neural dynamics of Bayesian Causal Inference in perception of audiovisual stimulus sequences.

## Results

During the EEG recording, we presented 23 human observers with sequences of auditory beeps and visual flashes in a four (1 to 4 flashes) × four (1 to 4 beeps) factorial design (Fig. 1). Participants estimated and reported either the number of flashes or the number of beeps. We combined a GLM-based and a Bayesian modelling analysis to characterize the computations and neural mechanisms of how the brain combines information from vision and audition to estimate the number of auditory and visual stimuli.

**Figure 1.**
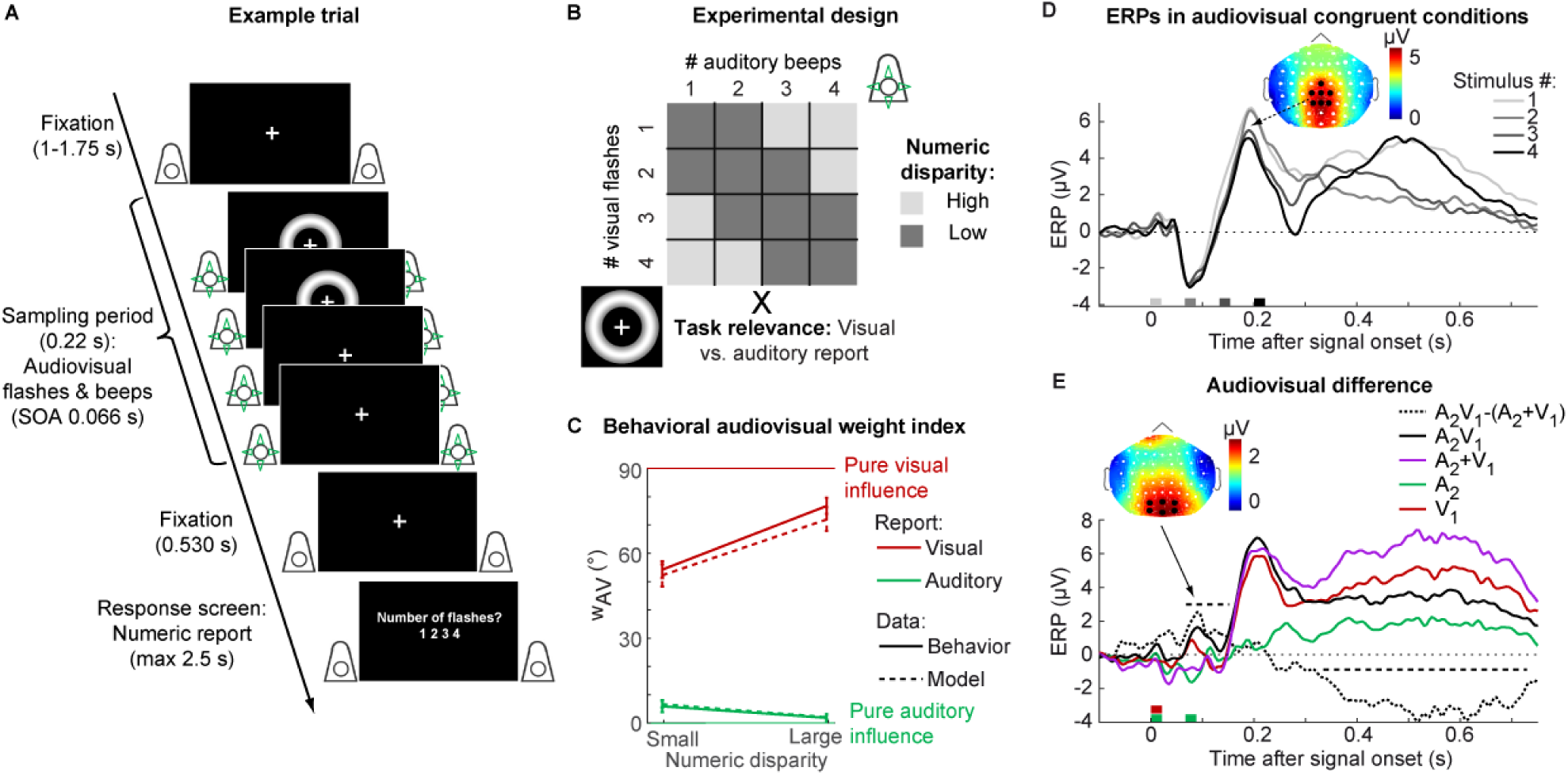
Example trial, experimental design and behavioral as well as EEG data. **(A)** Example trial of the flash-beep paradigm (e.g. two flashes and four beeps are shown) in which participants either report the number of flashes or beeps. **(B)** The experimental design factorially manipulated the number of beeps (i.e. one to four), number of flashes (i.e. one to four) and the task relevance of the sensory modality (report number of visual flashes vs. auditory beeps). To characterize the computational principles of the Bayesian Causal Inference (BCI) model, we reorganized these conditions into a two (task-relevance: auditory vs. visual report) × two (numeric disparity: high vs. low) factorial design for the GLM-based analysis of the behavioral and EEG data (e.g. audiovisual weight index). **(C)** The behavioral audiovisual weight index w_AV_ (across-participants circular mean and bootstrapped 68% CI; n=23) is shown as a function of numeric disparity (small: ≤ 1 vs. large: > 1) and task relevance (auditory vs. visual report). w_AV_ was computed for participants’ numeric reports (solid) and the BCI model’s predicted reports (dashed). w_AV_=90° for purely visual and w_AV_ = 0° for purely auditory influence. **(D)** Event-related potentials (ERPs; across-participants mean; n = 23) elicited by one to four stimuli in audiovisual congruent conditions averaged across parietal electrodes and the ERP topography at the peak (averaged over conditions with one to four stimuli). The x axis shows the stimulus onsets. **(E)** Difference between ERPs elicited by two beeps and one flash and the sum of the corresponding unisensory ERPs (i.e. V_1_A_2_-(A_2_+V_1_), black dotted), the unisensory auditory ‘two beeps’ (A_2_, green), visual ‘one flash’ (V_1_, red), audiovisual (A_2_V_1_, black solid) and the sum of the unisensory (A_2_+V_1_, pink) averaged across occipital electrodes. A positive component was significant from 65-150 ms (p = 0.040, two-sided cluster-based corrected randomization t_22_ test; see horizontal dashed line) and a negative component was significant 335-730 ms after stimulus onset (p < 0.001). The ERP topography at the peak of the positive component is shown.

### Behavior – Audiovisual weight index and Bayesian modelling

Using a general linear model (i.e. GLM, regression) approach, we computed a relative audiovisual weight index w_AV_ that quantifies the relative influence of the true number of beeps and flashes on participants’ numeric reports. The audiovisual weight index w_AV_ was analyzed as a function of numeric disparity between beeps and flashes (i.e. small ≤ 1 vs. large ≥ 2) x task-relevance (visual vs. auditory report). This audiovisual weight index ranges from pure visual (90°) to pure auditory (0°) influence. As shown in figure 1C and figure 2A, observers’ reported number of beeps was mainly influenced by the true number of beeps and only slightly – but significantly – biased by the true number of flashes (circular mean w_AV_ = 3.871, p < 0.001, one-sided randomization test on w_AV_ > 0°; i.e. a visual bias on auditory perception^18,19^). Conversely, the reported number of flashes was biased by the true number of beeps (circular mean w_AV_ = 65.483, p < 0.001, one-sided randomization test on w_AV_ < 90°), which is consistent with the well-known ‘sound induced flash illusion^10,19^. Yet, despite these significant biases operating from vision to audition and vice versa, observers did not fuse stimuli into one unified percept. Instead, the visual influence was stronger when the number of flashes was reported and the auditory influence was stronger when number of beeps was reported (effect of task on w__AV__, LRTS = 85.620, p < 0.001, randomization test of a likelihood ratio test statistic (LRTS); Table 1). As a result, observers reported different perceived numbers of flashes and beeps for audiovisual stimuli with a numeric disparity. Thus, participants flexibly adjusted the weights according to the task-relevant sensory modality. Crucially, this difference between auditory and visual report increased significantly for large relative to small numeric disparities. In other words, audiovisual integration broke down for large numeric disparities, when auditory and visual stimuli were more likely to be caused by independent sources (i.e. a significant interaction between task-relevance and numeric disparity, LRTS = 1.761, p < 0.001; for analysis of response times, see supplementary results and figure S1).

**Table 1.** Effects of numeric disparity (D) and task relevance (T) on the behavioral and the neural audiovisual weight index w_AV_ computed from numeric estimates decoded from EEG activity patterns (i.e. significant clusters).

| Effect | Behavioral $w_{AV}$ | | Neural $w_{AV}$ | | |
| --- | --- | --- | --- | --- | --- |
|  | LRTS | p | Cluster # | Time (s) | p |
| T | 85.620 | <0.001 | 1 | 0.42-0.54 | <0.001 |
|  |  |  | 2 | 0.62-0.66 | 0.007 |
| D | 1.624 | <0.001 | 1 | 0.40-0.48 | <0.001 |
| T X D | 1.761 | <0.001 | 1 | 0.56 | 0.039 |
|  |  |  | 2 | 0.68-0.72 | 0.001 |

Note: p values are based on randomization tests using a circular log-likelihood ratio statistic (LRTS). Effects for the neural $w_{AV}$ are cluster-based correction for the multiple time points. Only significant (i.e. $p < 0.05$ ) clusters are reported. $N = 23$ .

**Figure 2.**
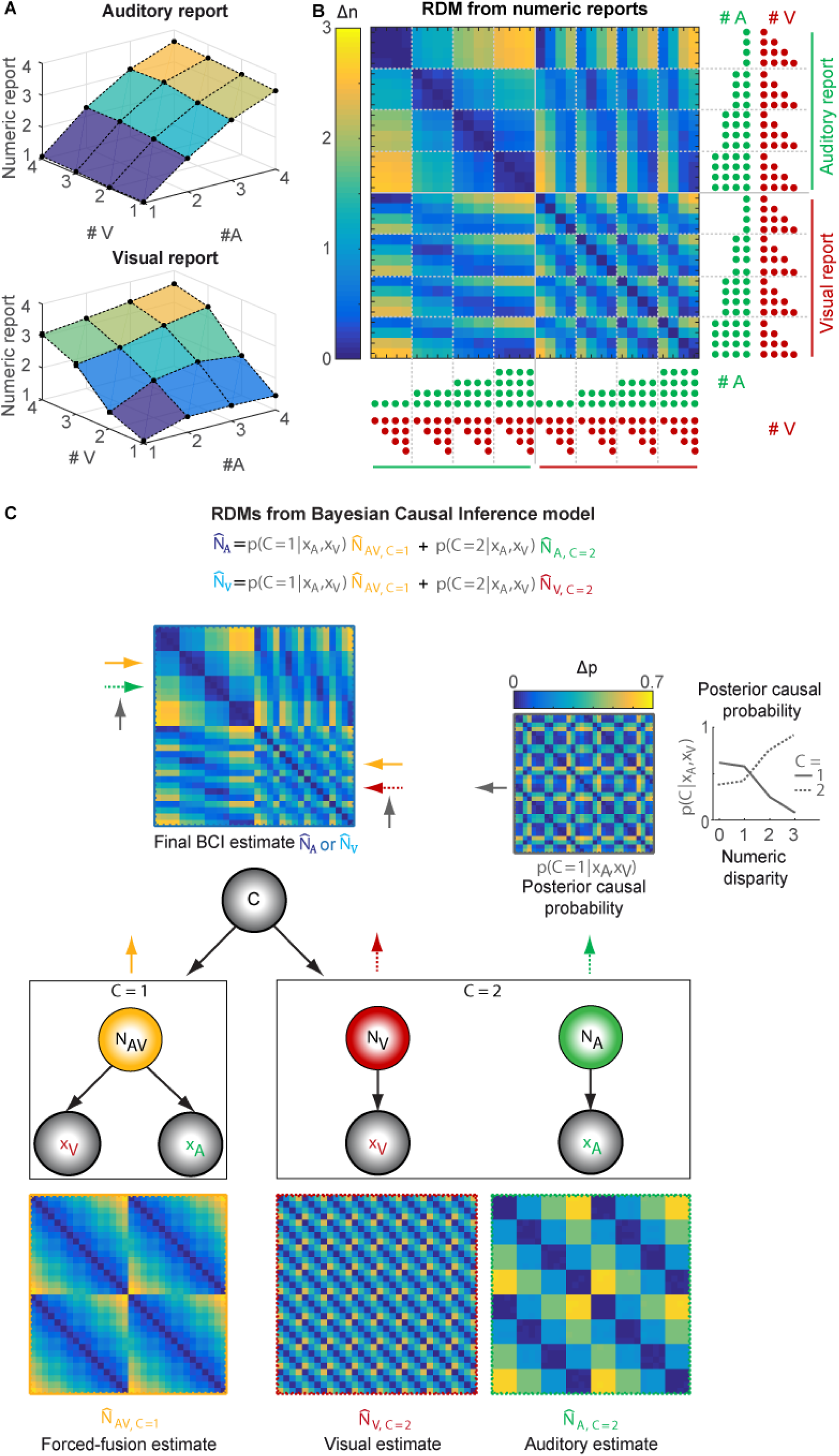
Representational dissimilarity matrices (RDMs) for numeric reports and estimates of the BCI model. **(A)** Participants’ auditory and visual numeric reports (across-participants’ mean; n = 23) are plotted as a function of the true number of visual (# V) and auditory (# A) stimuli, separately for auditory (top) and visual report (bottom). The auditory reports are more strongly influenced by the true number of auditory stimuli, while the visual reports are more strongly influenced the number of visual stimuli. Yet, crossmodal biases are also present. **(B)** RDM (across-participants’ mean) showing the absolute differences in participants’ numeric reports between all pairs of the 32 experimental conditions. The true number of beeps (green) and flashes (red) for each condition is indicated by the number of dots. **(C)** RDMs (across-participants’ mean) computed from the numeric estimates and the estimate of the posterior probability of a common cause of the BCI model: The generative BCI model assumes that in case of a common cause (C = 1), the “true” number of audiovisual stimuli N_AV_ is drawn from a common numeric prior distribution leading to noisy auditory (x_A_) and visual (x_V_) inputs. In case of independent causes (C = 2), the “true” auditory (N_A_) and visual (N_V_) numbers of stimuli are drawn independently from the numeric prior distribution. To account for the causal uncertainty, the final BCI estimate of the auditory or visual stimulus number (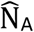 or 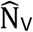, depending on whether auditory or visual modality is reported) is computed by combining the forced-fusion estimate of the auditory and visual stimuli 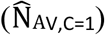 with the task-relevant unisensory visual (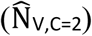) or auditory estimates 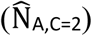, each weighted by the posterior probability of a common (C = 1) or independent (C = 2) causes, respectively (p(C| x_A_, x_V_)). Arrows indicate the influence of component estimates on the RDM of the final BCI estimate. The probability of independent causes (dashed) increases with larger numeric disparity and vice versa for the probability of common cause (n.b. they sum to unity).

Indeed, the model predictions in figure 1C show that this interaction between task-relevance and numeric disparity is a key feature of Bayesian Causal Inference. As this behavioral profile can be accounted for neither by the classical forced-fusion model that assumes audiovisual stimuli are fused into one single estimate (i.e. common source model) nor by the full-segregation model (i.e. independent source model), the Bayesian Causal Inference model was the winning model for explaining the behavioral data based on formal Bayesian model comparison (Table 2). Further, the decisional function ‘model averaging’ outperformed ‘model selection’ and ‘probability matching’ at the group level (see supplementary Table S1, consistent with^5^, but see ^20^). In the following, we will therefore focus selectively on Bayesian causal inference with model averaging.

**Table 2.** Results of the Bayesian model comparison of the Bayesian Causal Inference, a forced-fusion and a full-segregation model.

| | $p_{\text{common}}$ | $\mu_P$ | $\sigma_P$ | $\sigma_A$ | $\sigma_V$ | $R^2$ | relBIC | pEP | % win |
| --- | --- | --- | --- | --- | --- | --- | --- | --- | --- |
| Causal Inference<br>(model averaging) | 0.42±0.05 | 2.26±0.20 | 2.34±0.29 | 0.53±0.03 | 1.11±0.23 | 87.45±1.16 | 0 | 1 | 95.7 |
| Forced fusion | - | 2.10±0.22 | 4.38±0.60 | 1.17±0.04 | 1.45±0.04 | 61.68±1.58 | 8362.42 | 0 | 4.3 |
| Full segregation | - | 2.15±0.20 | 2.11±0.32 | 0.55±0.03 | 1.01±0.11 | 84.61±1.54 | 920.70 | 0 | 0 |

Note: $p_{\text{common}}$ , causal prior; $\mu_P$ , mean of the numeric prior; $\sigma_P$ , standard deviation of the numeric prior; $\sigma_A$ , standard deviation of the auditory likelihood; $\sigma_V$ , standard deviation of the visual likelihood; $R^2$ , coefficient of determination; relBIC, Bayesian information criterion at the group level, i.e. subject-specific BICs summed over all subjects ( $\text{BIC} = \text{LL} - 0.5 M \ln(N)$ , LL = log likelihood, M = number of parameters, N = number of data points) of a model relative to the Bayesian Causal Inference (“model averaging”) model (n.b. a smaller relBIC indicates that a model provides a better explanation of our data); pEP, protected exceedance probability, i.e. the probability that a given model is more likely than any other model, beyond differences due to chance). % win, percentage of participants in which a model won the within-participant model comparison based on BIC.

### EEG – conventional univariate ERP analysis

Event-related potentials (ERPs) revealed the typical sequence of ERP components in response to audiovisual flashes and beeps (Fig. 1D), i.e. P1 (∼ 50ms), N1 (100ms), P2 (200ms), N2 (280 ms) and P3 (> 300 ms)^21^. In line with previous studies^21-23^, we observed early multisensory interactions in the classical’ sound-induced flash illusion’ comparison (i.e. A__1__V_1_A_2_ vs. A_1_A_2_ + V_1_; Fig. 1E) over occipital electrodes starting at about ∼70 ms (i.e. measured from the onset of the first flash-beep slot). Further, we observed a negative audiovisual interaction component at 335-730 ms after stimulus onset. However, the current study did not focus on early multisensory interactions as evidenced in ERPs, but on the neural dynamics underlying Bayesian Causal Inference in perceptual decision-making.

### EEG – multivariate decoding and audiovisual weight index

To compute a neural audiovisual weight index w_AV_, we applied multivariate pattern analysis to single trial EEG activity patterns (i.e. 64 electrodes) of 20 ms time intervals. We trained a support-vector regression model on EEG activity patterns independently at each time point of the audiovisual congruent conditions to establish a mapping between EEG activity pattern and number of audiovisual stimuli. We then generalized to the congruent and incongruent conditions (i.e. leave-one-run out cross-validation). First, we ensured that we could decode the stimulus number for congruent trials significantly better than chance. Indeed, the decoder was able to discriminate between for instance three and four flash-beeps nearly immediately after the presentation of the fourth flash-beep (Fig. 3A) and thus before the ERP traces, when averaged over parietal electrodes, started to diverge (Fig. 1D). Pooling over all four congruent conditions, we observed better than chance decoding accuracy from around 100 ms to 740 ms measured from the onset of the first flash-beep slot (Fig. 3B).

**Figure 3.**
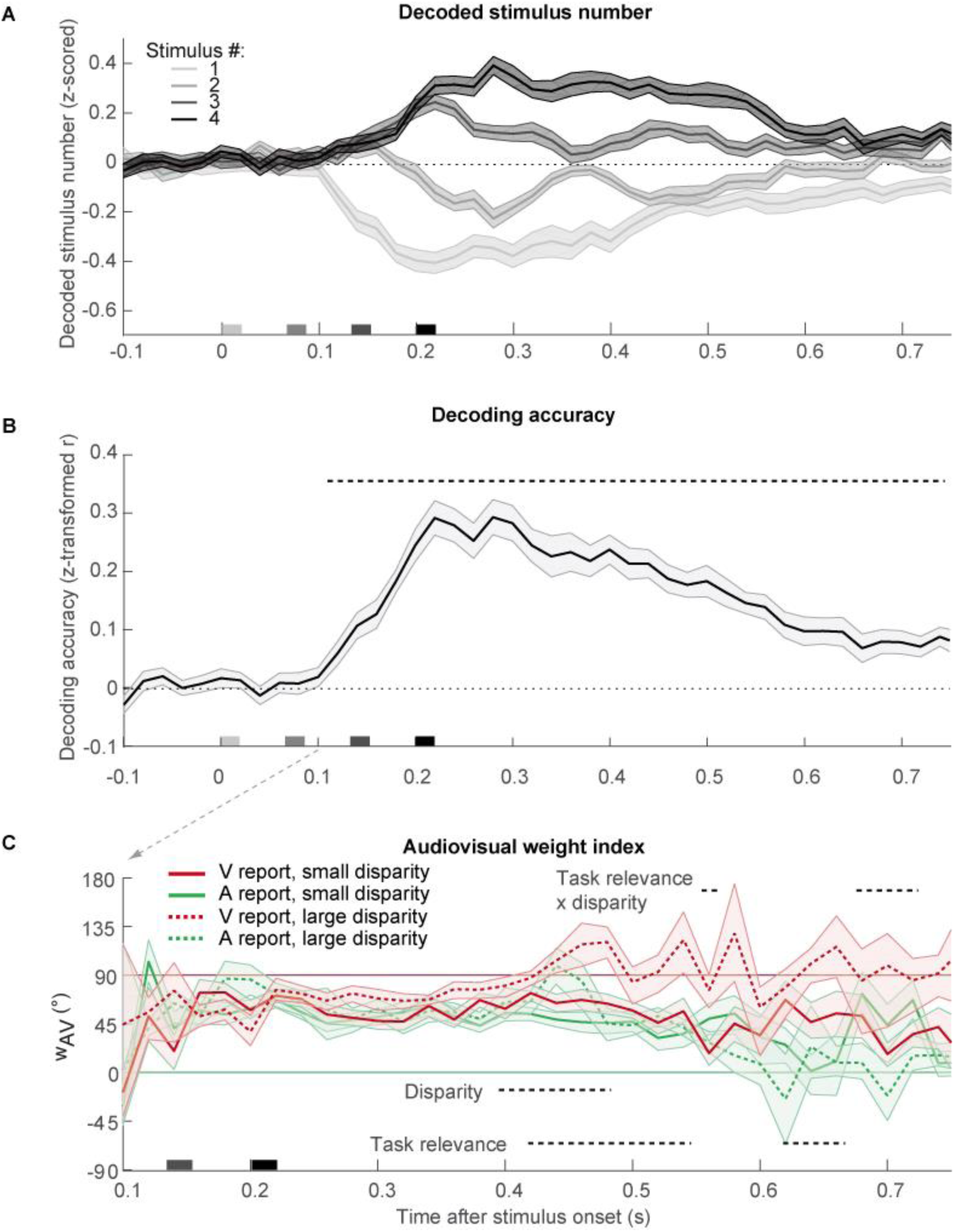
Decoded stimulus number, decoding accuracy, and relative audiovisual weight index. **(A)** Decoded stimulus number in audiovisual congruent conditions (across-participants mean ± SEM; n = 23). Note that the stimulus numbers {1, 2, 3, 4} as labels were transformed to {−1, −0.33, +0.33, +1} before decoding such that the decoder predicts a stimulus value of 0 in case of no information on stimulus number in EEG activity patterns) and decoded stimulus number was z scored in each run. Stimulus onsets are shown on the x axis. **(B)** Decoding accuracy (Fisher’s z-transformed correlation; across-participants mean ± SEM) of the decoders as a function of time. Decoding accuracy was computed as Pearson correlation coefficient between true and decoded stimulus number in audiovisual congruent conditions. Decoding accuracy was significant in a cluster from 120-740 ms (p < 0.001, one-sided cluster-based corrected randomization t_22_ test; see horizontal dashed line). **(C)** Neural audiovisual weight index w_AV_ (across-participants circular mean and bootstrapped 68% CI) as a function of numeric disparity (small vs. large), task relevance (visual (V) vs. auditory (A) report) and time. w_AV_ = 90° for purely visual and w_AV_ = 0° for purely auditory influence. Horizontal dashed lines indicate significant clusters for effects of numeric disparity, task relevance and their interaction (p < 0.05; cluster-based corrected randomization test based on a LRTS statistic, cf. Table 1).

We applied the same analysis approach as for behavioral responses to the audiovisual decoded numeric estimates and computed the neural audiovisual weight index w_AV_ which quantified the relative auditory and visual influences on the decoded number of flashes and beeps across poststimulus time (i.e. from 100 ms to 740 ms). We assessed how the neural audiovisual weight index was affected by numeric disparity between beeps and flashes (i.e. small ≤ 1 vs. large ≥ 2) and task-relevance (visual vs. auditory report) in a 2 × 2 repeated measures analysis (Fig. 3C and Table 1). We observed that the auditory influence was stronger for small relative to large numeric disparities from 400-480 ms poststimulus (i.e. effect of numeric disparity: 200-280 ms after the final flash-beep slot). Only when the numeric disparity was small and hence the two stimuli were likely to come from a common cause, did auditory stimuli impact the neural estimation of the number of flashes, which dominated the EEG activity patterns. Shortly later, i.e. 420-540 ms poststimulus, the influence of the auditory and visual stimuli on the decoded numeric estimate also depended on the sensory modality that needed to be reported (effect of task-relevance; for additional effects see Table 1). The number of flashes influenced the decoded numeric estimates more strongly for visual report, whereas the number of beeps influenced the decoded numeric estimates for auditory report. Crucially, at 560 ms and from 680 to720 ms poststimulus, we observed a significant interaction between task-relevance and numeric disparity, which is the key profile of Bayesian Causal Inference. As predicted by Bayesian Causal Inference (cf. Fig 1C), the audiovisual weight index for auditory and visual report were similar (i.e. integration) for small numeric disparity, but diverged (i.e. segregation) for large numeric disparities when it is unlikely that the flash and the beep sequences were generated by a common cause.

### Representational geometry of the numeric estimates of the Bayesian Causal Inference model and EEG activity pattern

Using representational dissimilarity analysis, we compared the representational geometry of the full-segregation auditory or visual, forced-fusion and the final Bayesian Causal Inference (BCI) estimates^24^ with the representational geometry of observers’ numeric reports (Fig. 2) and EEG activity patterns across poststimulus time (Fig. 4). First, we estimated the representational dissimilarity matrices (RDMs) by computing the pairwise absolute distance between the BCI model’s four numeric estimates, i.e. (i) the forced-fusion, the full-segregation (ii) auditory and (iii) visual and (iv) the final BCI estimates as well as the posterior causal probability across all 32 conditions. As shown in figure 2 C, the RDM for the forced-fusion estimate 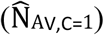 was a weighted average of the RDMs of full-segregation auditory 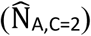 and visual 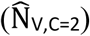 estimates. Further, because the auditory modality provides more precise temporal information (cf. Table 2) which is crucial for estimating the number of stimuli, the forced-fusion RDM is more similar to the auditory than the visual RDM. Finally, the RDM for the BCI estimate (i.e. 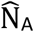 or 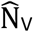, depending on the sensory modality that needs to be reported) combines the forced-fusion estimate 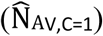 with the task-relevant unisensory visual 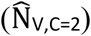 or auditory 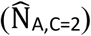 estimates (depending on report), weighted by the posterior probability of a common or separate causes, respectively (i.e. p(C = 1| x_A_, x_V_) or p(C = 2| x_A_, x_V_)). The probability of a common cause increased with smaller numeric disparity such that the influence of the forced-fusion estimate was greater for small numeric disparities. Figure 2B illustrates that the RDM computed from observers’ behavioral numeric reports was nearly identical to the BCI RDM. This match was confirmed statistically by a high correlation between the BCI (i.e. 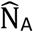 or 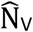) RDM and participants’ behavioral RDM (r = 0.878 ± 0.059, mean ± SEM, p < 0.001). Of course, this match between behavioral and BCI RDM was expected because the BCI RDM was computed from the predictions of the BCI model that well fit participants’ numeric reports (i.e. circular dependency; cf. Table 2).

**Figure 4.**
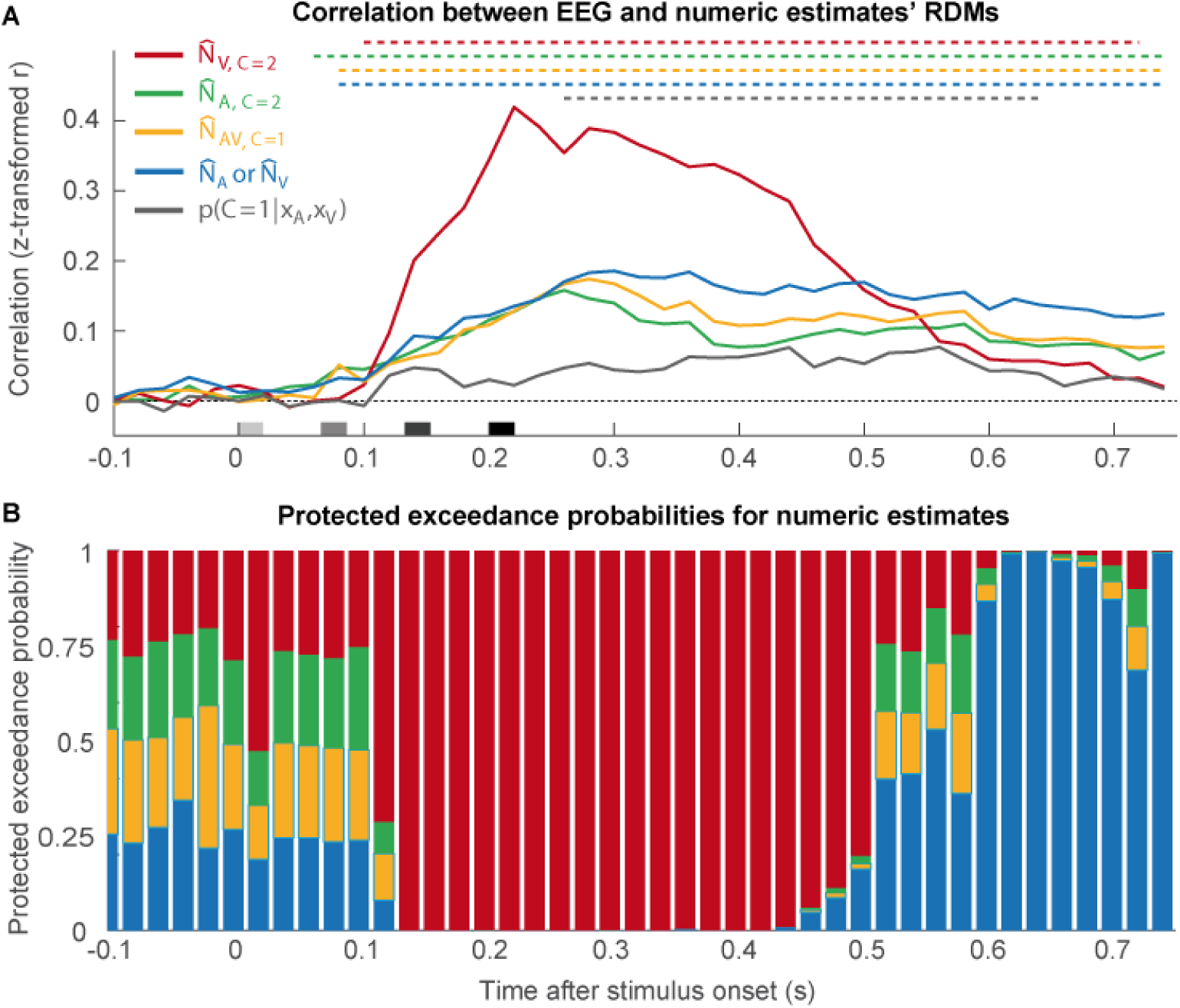
Correlation of the representational dissimilarity matrices of EEG activity patterns with the BCI model’s estimates and their exceedance probability. **(A)** Spearman’s rank correlation coefficients (across-participants mean of Fisher’s z-transformed r; n = 23) as a function of time. The correlation coefficients were computed between the RDMs of EEG activity patterns of all 64 electrodes and the RDMs of the BCI model’s internal estimates: i. the unisensory visual (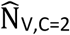, red), ii. the unisensory auditory (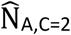, green) estimates under the assumption of independent causes (C = 2), iii. the forced-fusion estimate (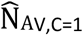, yellow) under the assumption of a common cause (C = 1) and iv. the final BCI estimate (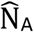 or 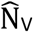, depending on the task-relevant sensory modality, blue) that averages the task-relevant unisensory and the reliability-weighted estimate by the posterior probability estimate of each causal structure (p(C = 1|x_A_, x_V_)). Color-coded horizontal dashed lines indicate clusters of significant correlation coefficients (p < 0.05; one-sided cluster-based corrected randomization t_22_ test). Stimulus onsets are shown along the x axis. **(B)** To identify which of the four numeric BCI estimates is most likely represented in the EEG activity patterns, we computed the protected exceedance probability (i.e. the probability that a given variable is more likely encoded in the EEG activity patterns than any other variable, beyond differences due to chance) for each of the four numeric BCI estimates as a function of time. The length of an estimate’s bar indicates the estimate’s protected exceedance probability (n.b.: the y axis indicates protected exceedance probabilities cumulated over the BCI estimates)‥

Next, we characterized the neural dynamics of Bayesian Causal Inference by comparing the representational geometry obtained from EEG activity patterns across time with the representational geometries of (i) the forced-fusion, the full-segregation (ii) auditory and (iii) visual and (iv) the final BCI estimates. As shown in figure 4A, the RDMs obtained from EEG activity patterns significantly correlated with the unisensory auditory RDM (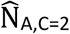; significant cluster 60-740 ms, p < 0.001, one-sided cluster-based corrected randomization t_22_ test), the unisensory visual RDM (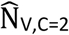; cluster 100-720 ms, p < 0.001), the forced-fusion RDM (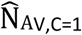; cluster 80-740 ms, p < 0.001) and the BCI RDM (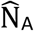 or 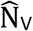; significant cluster 80-740 ms, p < 0.001). In short, the RDMs of EEG activity patterns correlated with multiple numeric estimates simultaneously. For the posterior probability of a common cause (p(C = 1|x_A_, x_V_), the correlation was weaker but significant in a later cluster (260-640 ms after stimulus onset, p < 0.001). The strong and sustained correlations of EEG RDMs and the RDMs of the four numeric estimates from the BCI model were expected because the four numeric estimates were highly correlated with one another. Hence, to account for these inherent correlations between these numeric estimates, we next computed the exceedance probability (i.e. the probability that the correlation with one numeric RDM was greater than that of any other RDMs) to determine which of the four numeric estimates was most strongly represented in the EEG activity patterns at a given time point (Fig. 4B). The exceedance probabilities showed that the EEG activity patterns predominantly encoded the unisensory visual estimate from 120 ms up to around 500 ms (i.e. 300 ms after the final flash-beep slot). This visual over auditory influence on EEG activity patterns at the scalp may be surprising, because the auditory sense exerts a stronger influence on observers’ reported numeric estimates (Fig. 1C) and provides more precise temporal information when estimated from observers’ numeric reports (cf. σ_A_ vs. σ_V_ in Table 2). Potentially, the visual neural sources elicit EEG activity patterns in sensor space that are more informative about the number of events (see methods section for caveats and critical discussion of the decoding analysis). Indeed, additional multivariate decoding analyses of the unisensory auditory and visual conditions showed that the number of visual stimuli could be more accurately decoded from visual EEG activity patterns than the number of auditory stimuli from auditory EEG activity patterns (Supplementary Fig. S2). Potentially, this advantage for visual decoding under unisensory stimulation may further increase in audiovisual context when the visual signal is task-relevant because of additional attentional amplification.

Crucially, from 450 ms poststimulus (i.e. 250 ms after the presentation of the final flash-beep; Fig. 4B), the EEG representational geometries progressively reflected the BCI estimate. Collectively, the model-based representational dissimilarity analysis suggests that Bayesian Causal Inference evolves by dynamic encoding of multiple sensory estimates. First, the EEG activity is dominated by the numeric unisensory and forced-fusion estimates (i.e. 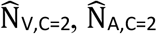, and 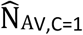) and later by the BCI estimate (i.e. 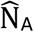 or 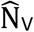) that takes into account the observers’ uncertainty about the world’s causal structure.

### EEG: Effect of prestimulus oscillations on the causal prior probability

Previous research demonstrated that observers perceived a sound-induced flash illusion more often for low prestimulus alpha power and/or high gamma and beta power over occipital (i.e. visual) cortices^16,17^. Within the framework of Bayesian Causal Inference, the occurrence of a sound-induced flash illusion may increase when visual precision is reduced or the causal prior (i.e. the probability of a common versus independent causes, also known as binding tendency^25^) is enhanced. We therefore investigated whether prestimulus oscillatory power (over occipital electrodes) alters participants’ multisensory perception as parameterized by the causal prior (p_common_) or the precision of visual representations (i.e. the reciprocal of σ_V_). For this, we sorted the trials into 10 deciles according to oscillatory power for each time and frequency point and re-fitted the causal prior or the precision of visual representations in the BCI model separately for each bin. Next, we computed the correlation of the causal prior or the visual precision with oscillatory power over deciles. This analysis showed that the causal prior correlated positively with gamma power (p = 0.036, two-sided cluster-based corrected randomization t_22_ test, starting at −220 ms prestimulus to stimulus onset) and negatively with alpha power (p = 0.042, from −320 ms to −100 ms prestimulus, Fig. 5A, B). Fig. 5C shows the weight index w_AV_ computed from participants’ behavior for each decile and the corresponding model predictions. Both human and model behavior showed more audiovisual influences (i.e,. w_AV_ indices shifted towards 0.5) for high gamma power and low alpha power. Crucially, these audiovisual biases operated from vision to audition and vice versa (i.e. a bidirectional bias which cannot be modelled by changes in visual precision). Hence, prestimulus gamma and alpha oscillations tune how the brain arbitrates between sensory integration and segregation. High gamma and low alpha power prior to stimulus presentation increase the brain’s tendency to bind stimuli across the senses. For completeness, we did not observe any significant effect of oscillatory power on the visual precision (Supplementary Fig. S3A).

**Figure 5.**
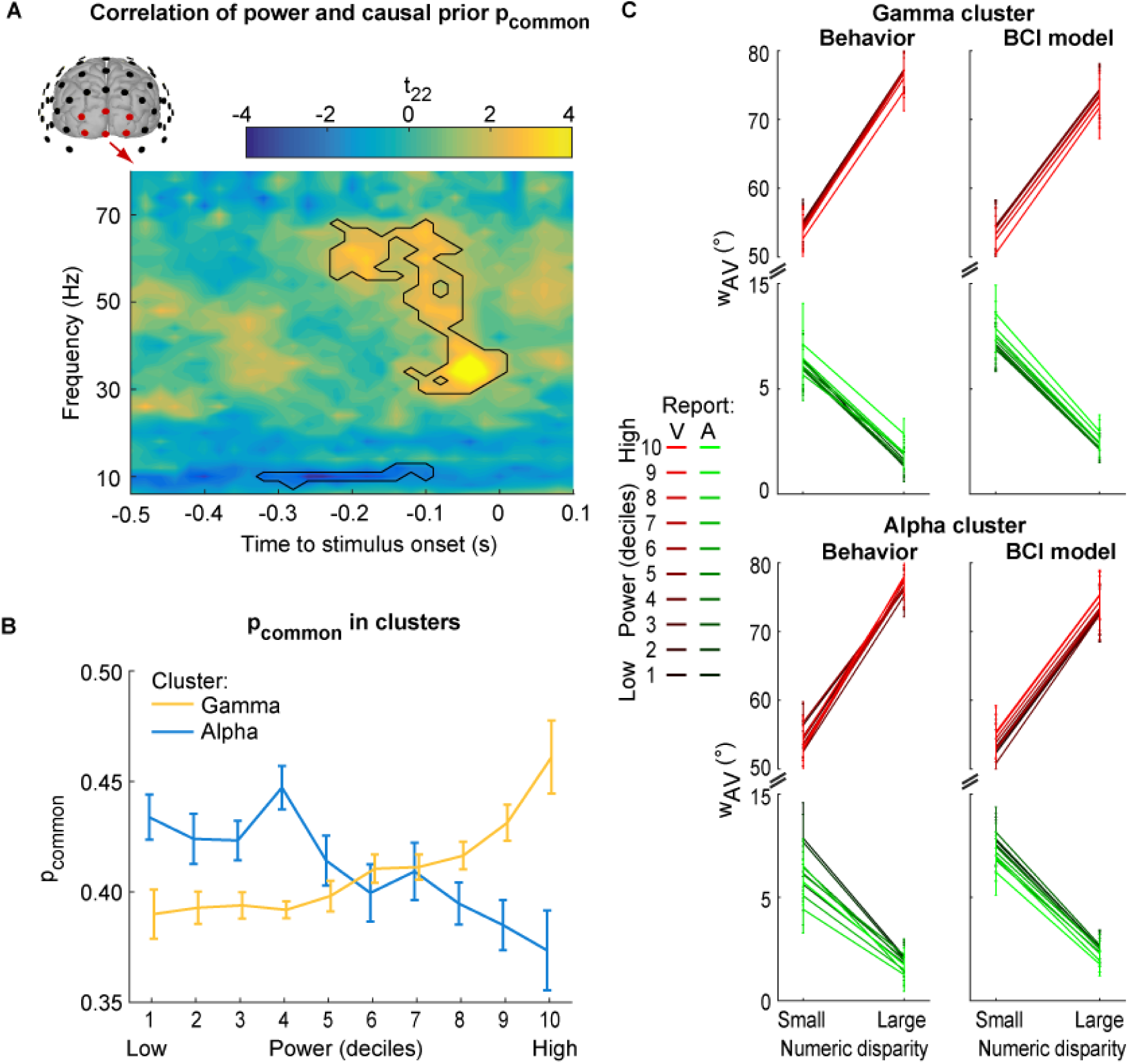
Effect of prestimulus oscillatory power on the BCI model’s causal prior (p_common_) over occipital electrodes. **(A)** Time-frequency t-value map (n = 23) for the correlation between p_common_ and the oscillatory power deciles averaged across occipital electrodes. Significant clusters (p < 0.05; two-sided cluster-based corrected randomization t_22_ test) are demarcated by a solid line. **(B)** p_common_ (across-participants’ mean ± within-participants’ SEM) as a function of power deciles averaged across the significant clusters in the gamma and alpha band shown in (A). **(C)** The weighting index w_AV_ (across-participants’ mean and bootstrapped 68% CI) computed from participants’ behavior (left column) and the BCI model’s predictions (right column) for each of the 10 power deciles shown in (B). w_AV_ is shown as a function of numeric disparity (small: ≤ 1 vs. large: > 1) and task relevance (A: auditory vs. V: visual report), separately for the significant cluster in the gamma (top) and alpha band (bottom). w_AV_ = 90° for purely visual and w_AV_ = 0° for purely auditory influence.

Given the prominent role of alpha oscillations in temporal binding^26,27^ in visual and multisensory perception, we next investigated whether the prestimulus alpha phase influenced the causal prior or visual precision. Using a similar sort-and-binning approach as for prestimulus power, we computed a circular-linear correlation between alpha phase and causal prior (or visual precision) over deciles as a function of prestimulus time. While there was again no significant effect of alpha phase on visual precision (Supplementary Fig. S3B), we observed a significant cluster from −160 ms to −80 ms prestimulus (p = 0.015, one-sided cluster-based corrected randomization t_22_ test), where alpha phase correlated significantly with participants’ causal prior (Fig. 6A): trials with a specific alpha phase led to a higher causal prior than trials with an opposing alpha phase. Importantly, the relation between alpha phase and causal prior progressed consistently over time at alpha frequency (i.e. 10 Hz; Fig. 6C). In support of this, a sinusoidal model in which the phase of an alpha oscillation modulated the causal prior outperformed a model that did not include a sinusoidal modulation from −280 ms to −80 ms prestimulus in 20 out of 23 participants (individual F_2,107_ tests, p < 0.05; see Supplementary Fig. S4 for individual data from four representative participants). However, the relation of alpha phase and causal prior was not consistent across participants (z_22_ = 2.486, p = 0.082, Raleigh test, Fig. 6B). These differences between participants are expected and may arise from differences in cortical folding and hence orientations of the underlying neural sources. To account for these differences across participants, we therefore aligned the alpha phase individually for each participant, such that the phase at the peak group effect at −160 ms prestimulus was consistent across participants (cf. Supplementary Fig. S5 for data without phase-alignment). Figure 6C and D show that the alpha phase modulates the causal prior across nearly three cycles which is consistent across participants. Collectively, these results demonstrate that the power and phase of prestimulus alpha oscillations influence observers’ causal prior, which formally quantifies their apriori tendency to bind signals from audition and vision into a coherent percept.

**Figure 6.**
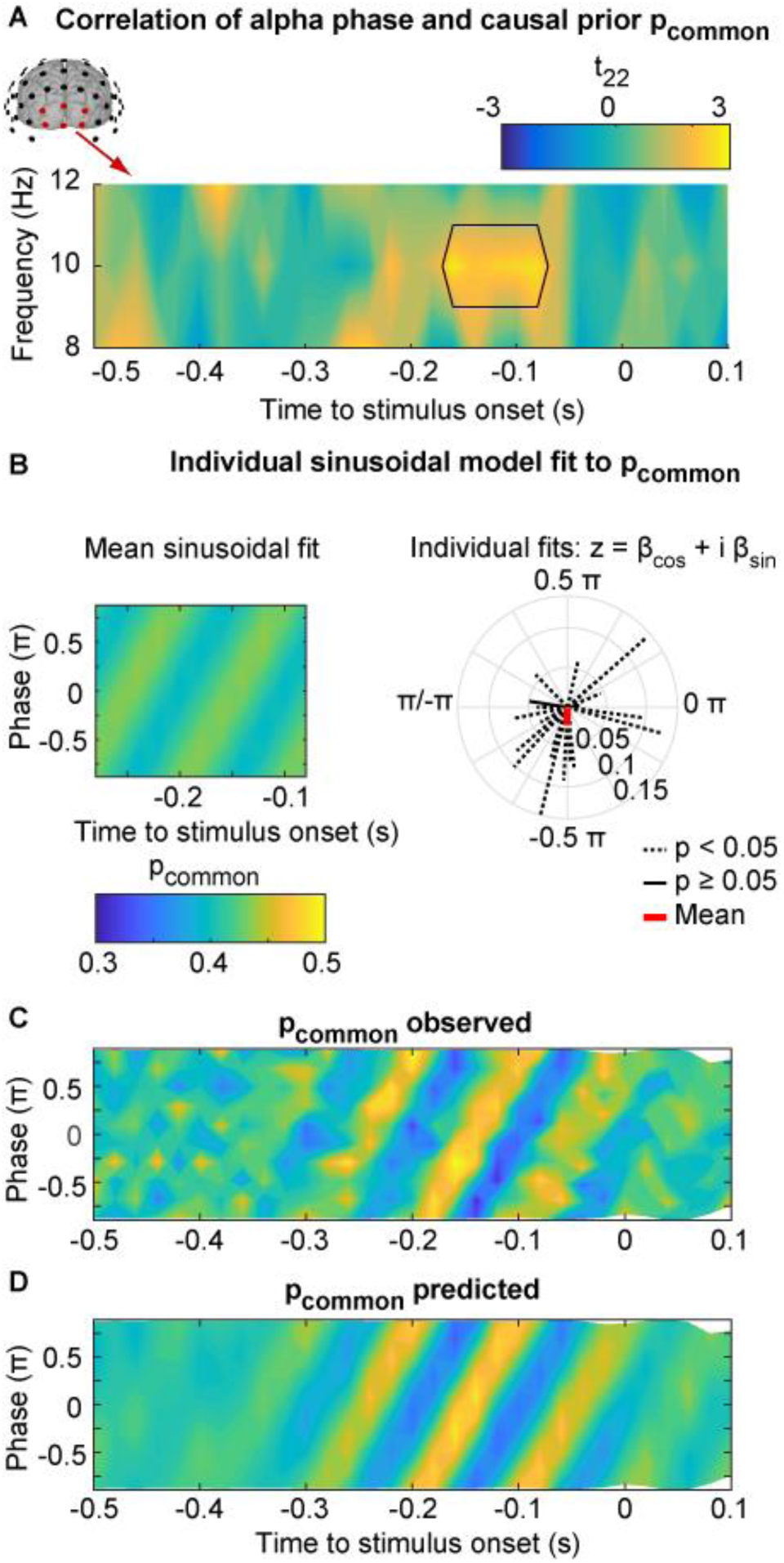
Effect of prestimulus alpha phase on the BCI model’s causal prior (p_common_) in occipital electrodes. **(A)** Time-frequency t-value map (n = 23) for the circular-linear correlation between p_common_ and the phase deciles of alpha oscillations for 8, 10 and 12 Hz and averaged across occipital electrodes. Significant clusters (p < 0.05; one-sided cluster-based corrected randomization t_22_ test) are demarcated by a solid line. **(B)** Left: Across participants’ mean predicted p_common_ (colour coded) as a function of time (x-axis) and 10 Hz alpha phase bin (y-axis, alpha phase indicated as fraction of π). The predicted p_common_ is based on the constrained sinusoidal model which predicts p_common_ for a particular decile and time by a single sine and cosine of the phase at 10 Hz averaged across trials in this particular decile and time bin (see eq. 9). Hence, this constrained sinusoidal model assumes that the modulation of p_common_ by alpha phase evolves at 10 Hz over time. Right: Individual model fits are significant (p < 0.05; individual F_2,107_ tests, dashed lines) in 20 of 23 participants, but the sinusoidal model’s phase angles (i.e. Φ_*subject*_ = angle(z) = angle(βcos + i βsin)) do not deviate significantly from a circular uniform distribution (z_22_ = 2.486, p = 0.082, Raleigh test) across participants. **(C,D)**. Across-participants’ mean observed (C) and predicted (D) p_common_ is shown coded in color as a function of time (x-axis) and 10 Hz alpha phase bin (y-axis, alpha phase indicated as fraction of π). The phases of the observed and predicted p_common_ were aligned across participants (at the peak of the non-aligned effect with alpha phase = −0.29 π at −160 ms; see methods section and Supplementary Fig. S3B) before averaging across participants. The predicted p_common_ (D) is based on the sinusoidal models that predicted p_common_ by alpha phase across the deciles independently for each time point (i.e. time-specific alpha phase models) and participant (i.e. eq. 8).

### EEG: The relationship of prior stimulus history, prestimulus alpha power and the causal prior probability

Previous research has shown that prior stimulus history influences observers’ binding tendency^28-30^. For instance, prior congruent audiovisual speech stimuli increased observers’ tendency to bind incongruent audiovisual signals into illusionary McGurk percepts^28^. Hence, we investigated whether the numeric disparity of previous flash-beep stimuli (going back in history to five trials prior to stimulus onset) influenced observers’ causal prior on the current trial. Indeed, as shown in figure 7A, a 2 (numeric disparity: small vs. large) × 5 (stimulus order: 1, 2, 3, 4, 5 trials back) repeated measures ANOVA revealed a significant main effect of numeric disparity (F_1,21_ = 6.260, p = 0.021, partial η^2^ = 0.230) and a significant interaction between numeric disparity and stimulus order (F_2.6,54.9_ = 4.060, p = 0.015, partial η^2^ = 0.162). Post-hoc tests for the effect of numeric disparity separately for specific stimulus order showed that the effect of numeric disparity was most pronounced for the first-and second-order previous stimulus (first order: t_22_ = 3.731, p = 0.001, Cohen’s d = 0.778; marginally significant second order: t_22_ = 2.042, p = 0.053, Cohen’s d = 0.426; two-sided paired t tests) and tapered off with stimulus order.

**Figure 7.**
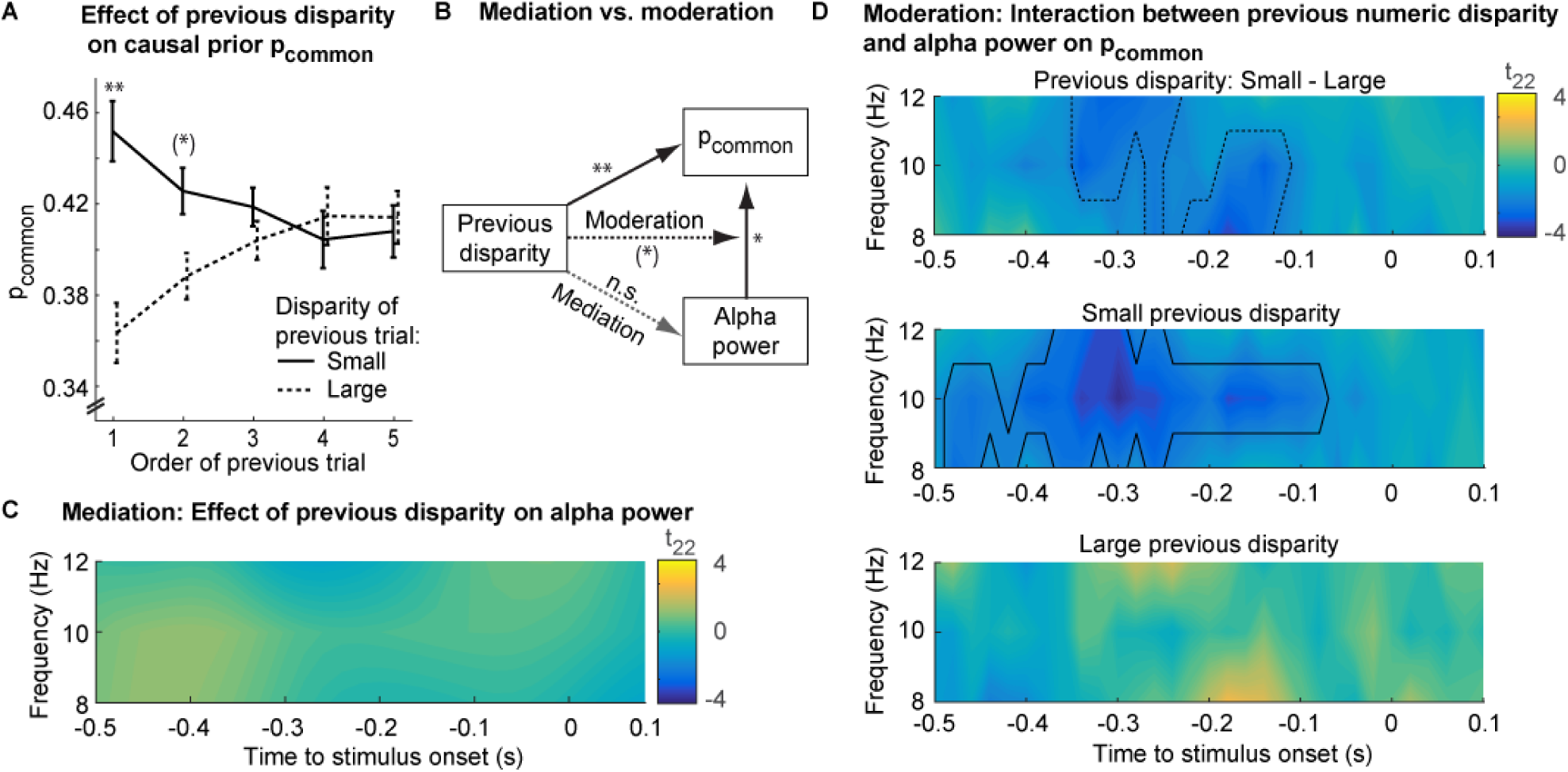
The effect of prior numeric disparity on the BCI model’s causal prior (p_common_) and prestimulus alpha power. **(A).** Effect of previous numeric disparity on causal prior p_common_ (across-participants’ mean ± within-participants’ SEM, n = 23) as a function of the numeric disparity (small: ≤ 1 vs. large: > 1) of the previous trial of order 1-5. The effect of previous numeric disparity on causal prior decays with increasing trial order Asterisks denote statistical significance in two-sided one-sample t_22_ tests (**: p < 0.01, *: p < 0.05, (*): p < 0.1, n.s.: not significant). **(B)** Mediation and moderation: Previous numeric disparity and pre-stimulus alpha power significantly predict p_common_. However, previous numeric disparity does not predict pre-stimulus alpha power. Hence, pre-stimulus alpha power does not mediate the effect of numeric disparity on causal prior. Instead, we observed a marginally significant interaction between previous numeric disparity and alpha power suggesting that previous numeric disparity moderates the effect of pre-stimulus alpha power on p_common_. **(C)** Mediation: Effect of previous numeric disparity on alpha power: Time-frequency t-value map (n = 23, averaged over occipital electrodes) reveals no significant difference in prestimulus alpha power between small versus large previous numeric disparity (i.e. trial order 1). All clusters p > 0.05 (two-sided cluster-based corrected randomization t_22_ test). **(D)** Moderation: Interaction effect between previous numeric disparity and alpha power on causal prior: Time-frequency t-value maps (n = 23) for the correlation between p_common_ and the alpha power averaged across occipital electrodes over deciles for trials with large (lower panel) or small (middle panel) numeric disparity of the previous trial and the difference in correlations between small and large disparity of the previous trial (top panel). We observed only a marginally significant interaction between previous disparity and alpha power on p_common_ (p < 0.1; two-sided cluster-based corrected randomization t_22_ test demarcated by dashed line in top panel). Prestimulus alpha power correlated significantly with observers’ causal prior only if the previous trial was of a small numeric disparity (p < 0.05; demarcated by a solid line, middle panel).

Our results so far suggest that previous stimulus history (i.e. numeric disparity of previous trials) and pre-stimulus alpha power predict observers’ tendency to bind audiovisual signals. This raises the intriguing question whether the effect of previous stimulus history is mediated by alpha power. For instance, given the well-established role of pre-stimulus alpha oscillations in visual perception^31-36^ and attention^37^, one may argue that alpha power is adjusted according to observers’ causal expectations based on prior stimulus history. Contrary to this conjecture, the numeric disparity of previous stimuli did not significantly predict alpha power (Fig. 7C; all clusters p > 0.05; Bayes factors provided substantial evidence in favour of a null effect, supplementary Fig. S8). However, we observed a marginally significant interaction between numeric disparity of the previous trial and alpha power on observers’ causal prior in two clusters from −340 to −240 ms, (p = 0.069) and from −220 to −120 ms (p = 0.096; two-sided cluster-based corrected randomization t_22_ test, Fig. 7C, top panel). The correlation between alpha power and the observers’ causal prior was prominent when prior numeric disparity was small (cluster from −480 to −80 ms, p = 0.006), but not significant when previous numeric disparity was large (i.e. all clusters p > 0.05). In summary, alpha power did not mediate, but to some extent (i.e. only marginally significant) moderated the effect of stimulus history on observers’ causal prior, i.e. their tendency to bind audiovisual signals (Fig. 7B).

## Discussion

To form a coherent percept of the world, the human brain needs to integrate signals arising from a common cause, but segregate signals from independent causes. Perception thus relies crucially on inferring the world’s causal structure^1,2^. To characterize the neural dynamics of how the brain solves this binding problem, we presented participants with sequences of beeps and flashes that varied in their numeric disparity.

Behaviorally, the number of beeps biased observers’ perceived number of flashes – a phenomenon coined sound-induced flash illusion^10^. Conversely, the number of flashes biased observers’ perceived number of beeps^18,19^, albeit only to a small degree. This asymmetry of crossmodal biases operating from vision to audition and vice versa can be attributed to the smaller precision of vision for temporal estimation, which is consistent with forced-fusion models of reliability-weighted integration^3,4,9^ (and Bayesian Causal Inference models, cf. Table 2). Crucially, as predicted by Bayesian Causal Inference, participants did not fully fuse auditory and visual stimuli into one unified percept, but they reported different numeric estimates for the flash and beep components of numerically disparate flash-beep stimuli. Moreover, audiovisual integration and crossmodal biases decreased for large numeric disparities, when the flash and beep sequences were unlikely to arise from a common cause^5,11^. Thus, observers flexibly arbitrated between audiovisual integration and segregation depending on the probabilities of the underlying causal structures as predicted by Bayesian Causal Inference (see Fig 2C).

At the neural level, our univariate and multivariate EEG analyses revealed that the computations and neural processes of multisensory interactions and Bayesian Causal Inference dynamically evolve poststimulus. Initially, the univariate ERP analyses revealed an early audiovisual interaction effect starting at about 70 ms poststimulus that is related to the visual P1 component and has previously been shown to be susceptible to attention^23^. Potentially, these early non-specific audiovisual interactions enhance the excitability in visual cortices and the salience of the visual input and may thereby facilitate the emergence of the sound-induced flash illusion^38^. Our multivariate EEG analyses revealed that the audiovisual weight index w_AV_ was influenced by both auditory and visual inputs until 400 ms postimulus, though with a slightly stronger influence of the visual input. This visual dominance in the multivariate pattern decoding may at least partly explain the surprisingly strong correlation between EEG activity pattern and the unisensory visual segregation estimate in the RDM analysis reaching a plateau from 200 ms to 400 ms poststimulus (see methods section for further discussion about methodological caveats). In addition, the posterior probability over causal structures is decodable from EEG activity patterns shortly after the final flash-beep slot. Likewise, the weight index w_AV_ indicated an early numeric disparity effect at about 400 ms poststimulus (i.e. 200 ms after the final stimulus slot; Fig 3). Thus, causal inference starts immediately after stimulus presentation based on numeric disparity and influences early audiovisual interactions and biases as quantified by the neural weight index. However, only relatively late, starting at about 200-300 ms and peaking at 400 ms after the onset of the final stimulus slot, does the brain compute numeric estimates consistent with Bayesian Causal Inference by averaging the forced-fusion estimate with the task-relevant unisensory estimate weighted by the posterior probabilities of common and independent causal structures (i.e. model averaging). The exceedance probability of the hierarchical Bayesian Causal Inference estimate steadily rises until its peak, where it outperforms all other numeric estimates in accounting for the representational geometries obtained from EEG activity patterns (i.e. exceedance probability ≈ 1). Likewise, the relative audiovisual weight index w_AV_ revealed a task-relevance by numeric-disparity interaction at similar latencies as the characteristic qualitative profile for Bayesian Causal Inference.

This dynamic evolution of neural representations dovetails nicely with a hierarchical organization of Bayesian Causal Inference that has recently been shown in fMRI research^12,13^: Low-level sensory areas represented sensory estimates mainly under the assumption of separate causes, whereas posterior parietal areas integrated the signals weighted by their sensory precision under the assumption of a common cause. Only at the top of the cortical hierarchy, in anterior parietal areas, did the brain form a final Bayesian Causal Inference estimate that takes into account the observers’ uncertainty about the signals’ causal structure. Collectively, fMRI and EEG research jointly suggest that computations involving unisensory estimates rely on lower-level regions at earlier latencies, while Bayesian Causal Inference estimates that take into account the world’s causal structure arise later in higher-level cortical regions. Previous fMRI research implicated prefrontal cortices in the computations of the causal structure^28,39^, which may in turn inform the integration processes in parietal and temporal cortices^40^.

A recent neural network model with a feedforward architecture by Cuppini et al. (2017, 2014)^41,42^ suggests that this explicit causal inference relies on a higher convergence layer, while the audiovisual biases in numeric estimates may be mediated via direct connectivity between auditory and visual layers and emerge from spatiotemporal receptive fields in auditory and visual processing. In contrast to such a feed-forward architecture, we generally observed a mixture of multiple representations that were concurrently expressed in EEG activity patterns, even though different numeric estimates dominated neural processing at different poststimulus latencies. Therefore, we suggest that Bayesian Causal Inference is iteratively computed via multiple feed-back loops across the cortical hierarchy whereby numeric estimates as well as causal inferences are recurrently updated as the brain accumulates knowledge about the causal structure and sources in the environment^14,15^.

In Bayesian inference, prior knowledge and expectations are crucial to guide the perceptual interpretation of the noisy sensory inputs^43^. Multisensory perception in particular relies on a so-called causal prior that quantifies observers’ prior beliefs about the world’s causal structure^1,2^. A ‘high’ causal prior (i.e. the belief that signals come from a common cause) influences multisensory perception by increasing observers’ tendency to bind audiovisual signals irrespective of the signals’ instantaneous intersensory congruency^28^. In the current study, we investigated whether the neural activity prior to stimulus onset is related to observers’ causal prior. Indeed, low prestimulus alpha power and high gamma power were associated with a high causal prior, i.e. they increased participants’ tendency to integrate audiovisual stimuli. Accumulating research has shown effects of prestimulus alpha power on perceptual decisions such as detection threshold, decisional biases or perceptual awareness^31-36^. Further, low alpha power was also shown to increase the occurrence of the sound-induced flash illusion^16,17,26^ (though see Keil et al.^16^ for an effect in beta power). In our study, low prestimulus alpha power predicted a larger causal prior leading to stronger bidirectional interactions between audition and vision and audiovisual biases (see figure 5C, audiovisual weight index w_AV_). These enhanced audiovisual interactions might be explained by a tonic increase in cortical excitability for states of low alpha oscillatory power and associated high gamma power (though see Yuval-Greenberg et al.^44^ for a cautionary note). Moreover, if peaks and troughs of alpha oscillatory activity are modulated asymmetrically^45^, low alpha power may also induce larger alpha troughs, thereby extending the temporal windows where gamma bursts and audiovisual interactions can occur^46,47^. Indeed, our results show that observers’ causal prior depends not only on the tonic level of alpha power, but also on its phase. Prestimulus alpha phase may thus influence audiovisual binding by defining the optimal time window in which neural processing can interact across auditory, visual and association areas, thereby modulating the temporal parsing of audiovisual signals into one unified percept^27,48,49^.

Next we investigated whether the fluctuations in alpha power may enable observers to adapt dynamically to the statistical structure of the sensory inputs. Previous research has shown that prior exposure to congruent signals increases observers’ tendency to integrate sensory signals, while exposure to incongruent signals enhances their tendency to process signals independently (^28,29^, but see^30^). In the current study, we also observed that previous low numeric disparity trials predicted a greater causal prior or tendency to bind audiovisual signals into a coherent percept. Surprisingly, however, the numeric disparity of previous audiovisual stimuli did not significantly influence alpha power. It only modulated the effect of alpha power on the causal prior (i.e. a marginally significant interaction between alpha power and prior numeric disparity). More specifically, alpha power correlated with observers’ causal prior mainly when previous stimuli were of low rather than large numeric disparity.

Collectively, our results show that observers’ causal prior dynamically adapts to the statistical structure of the world (i.e. previous audiovisual numeric disparity), but that these adaptation processes are not mediated by fluctuations in alpha power. Instead, spontaneous (i.e. as yet unexplained by stimulus history) fluctuations in prestimulus gamma and alpha power as well as alpha phase correlated with observers’ causal prior. Alpha power, phase and frequency (i.e. speed)^50-52^ together with gamma power may thus dynamically set the functional neural system into states that facilitate or inhibit interactions across brain regions^53,54^ and temporal parsing of audiovisual signals into common percepts^27,47^.

In conclusion, to our knowledge this is the first study that resolves the neural computations of hierarchical Bayesian Causal Inference in time. We show that pre-stimulus oscillatory alpha power and phase correlates with the brain’s causal prior as a binding tendency that guides how the brain dynamically arbitrates between sensory integration and segregation (see ^55-58^ for related studies showing that top-down predictions may be furnished via alpha/beta oscillations). Initially, about 70 ms after stimulus presentation, we observed non-specific audiovisual interactions, which may increase the bottom-up salience of sensory signals. Our multivariate analyses suggested that unisensory numeric estimates initially dominated the EEG activity pattern. Only later, from about 200-400 ms after the final stimulus slot, EEG signals encoded the Bayesian Causal Inference estimates that combine the forced-fusion and task-relevant segregation estimates weighted by the probabilities of common and independent cause models (i.e. model averaging). Thus, consistent with the notion of predictive coding, the brain may accumulate evidence concurrently about i. auditory (or visual) numeric estimates and ii. the underlying causal structure (i.e. whether auditory and visual signals come from common or independent sources) over several hundred milliseconds via recurrent message passing across the cortical hierarchy to compute Bayesian Causal Inference estimates^15,59^. By resolving the computational operations of multisensory interactions in human neocortex in time, our study reveals the hierarchical nature of multisensory perception. It shows that the brain dynamically encodes and re-updates computational priors and multiple numeric estimates to perform hierarchical Bayesian Causal Inference.

## Methods

### Participants

After giving written informed consent, 24 healthy volunteers participated in the EEG study based on previous calculations of statistical power. One participant did not attend the interview session and was excluded. Thus, data from 23 participants were analyzed (10 female; mean age 36.0 years, range 25-61 years). Participants were screened for current or former psychiatric disorders (as verified by the screening questions of the structured clinical interview for DSM IV axis I disorders, SCID-I, German version), cardio-vascular disorders, diabetes and neurological disorders. One participant reported an asymptomatic arteriovenous malformation. Because behavioral and EEG was inconspicuous, the participant was included. All participants had normal or corrected-to-normal vision and audition. The study was approved by the human research review committee of the Medical Faculty of the University of Tuebingen at the University Hospital Tuebingen (approval number 728/2014BO2).

### Stimuli

The flash-beep paradigm was an adaptation of previous “sound-induced flash illusion” paradigms^7,10^. The visual flash was a circle presented in the centre of the screen on a black background (i.e. 100% contrast; Fig. 1A) briefly for one frame (i.e, 16.7 ms, as defined by the monitor refresh rate of 60 Hz). The maximum grayscale value (i.e. white) of the circle was at radius 4.5° with smoothed inner and outer borders by defining the grayscale values of circles of smaller and larger radius by a Gaussian of 0.9° STD visual angle. The auditory beep was a pure tone (2000 Hz; ∼ 70 dBSPL) of 27 ms duration including a 10 ms linear on/off ramp. Multiple visual flashes and auditory beeps were presented sequentially at a fixed SOA of 66.6 ms (see below).

### Experimental design

In the flash-beep paradigm, participants were presented with a sequence of i. one, two, three or four flashes and ii. one, two, three or four beeps (Fig. 1A). On each trial, the number of flashes and beeps were independently sampled from one to four leading to four levels of numeric audiovisual disparities (i.e. zero = congruent to four = maximal level of disparity; Fig. 1B). Each flash and/or beep were presented sequentially in fixed temporal slots that started at 0, 66.7, 133, 200 ms. The temporal slots were filled up sequentially. For instance, if the number of beeps was three, they were presented at 0, 66.6, 133 and 200 ms, while the fourth slot was left empty. Hence, if the same number of flashes and beeps were presented on a particular trial, beeps and flashes were presented in synchrony. On numerically disparate trials, the ‘surplus’ beeps (or flashes) were added in the subsequent fixed time slots (e.g. in case of 2 flashes and 3 beeps: we present 2 flash-beeps at 0 and 66.6 ms in synchrony and a single beep at 133 ms).

Across experimental runs, we instructed participants to selectively report either the number of flashes or beeps and to ignore the stimuli in the task-irrelevant modality. Hence, the 4 × 4 × 2 factorial design manipulated (i) the number of visual flashes (i.e. one, two, three or four), (ii) the number of auditory beeps (i.e. one, two, three or four) and (iii) the task relevance (auditory-vs. visual-selective report) yielding 32 conditions in total (Fig. 1B). For analyses, we reorganized trials based on their absolute numeric disparity (|#A - #V| ≤ 1: small numeric disparity; |#A - #V| > 1: large numeric disparity). Thus, we analyzed the data in a 2 (task relevance: visual vs. auditory report) × 2 (numeric disparity) factorial design.

The duration of a flash-beep sequence was determined by the number of sequentially presented flash and/or beep stimuli (see above for the definition of temporal slots). Irrespective of the number of flashes and/or beeps, a response screen was presented 750 ms after the onset of the first flash and beep for a maximum duration of 2.5 s instructing participants to report their perceived number of flashes (or beeps) as accurately as possible by pushing one of four buttons. The order of buttons was counterbalanced across runs to decorrelate motor responses from numeric reports. On half of the runs, the buttons from left to right corresponded to one to four number of stimuli; on the other half they corresponded to four to one. After a participant’s response, the next trial started after an inter-trial interval of 1-1.75 s.

In every experimental run, each of the 16 conditions was presented 10 times. Participants completed 4 runs of auditory-and 4 runs of visual-selective report in a counterbalanced fashion (except for one participant performing 5 runs of auditory and 3 of visual report). Further, each participant completed two unisensory runs with visual or auditory stimuli only (i.e. 4 unisensory conditions presented 40 times per run) from which we computed the difference wave (see below). Before the actual experiment, participants completed 56 practice trials.

### Experimental setup

Psychtoolbox 3.09 (www.psychtoolbox.org) ^60,61^ running under MATLAB R2016a (MathWorks) presented audiovisual stimuli and sent trigger pulses to the EEG recording system. Auditory stimuli were presented at ≈ 70 dB SPL via two loudspeakers (Logitech Z130) positioned on each side of the monitor. Visual stimuli were presented on an LCD screen with a 60 Hz refresh rate (EIZO FlexScan S2202W). Button presses were recorded using a standard keyboard. Participants were seated in front of the monitor and loudspeakers at a distance of 85 cm in an electrically shielded, sound-attenuated room.

### Behavioral Analysis

#### Overview of general linear model and Bayesian modelling analysis for behavioral data

To characterize how human observers arbitrate between sensory integration and segregation, we developed a general linear model (GLM)-based and a Bayesian modelling analysis approach.

The GLM-based analysis computed a relative weight index w_AV_ which quantified the relative influence of the auditory and the visual numeric stimuli on observers’ auditory and visual behavioral numeric reports. This GLM-based analysis allowed us to reveal audiovisual weight profiles in our 2 (numeric disparity) × 2 (task-relevance) factorial design that are qualitatively in line with the principles of Bayesian Causal Inference (Fig. 1C).

The Bayesian modelling analysis fitted the full-segregation, the forced-fusion and the Bayesian Causal Inference (BCI) model to the behavioral numeric reports with different decision functions. We then used Bayesian model comparison to determine the model that is the best explanation for observers’ behavioral data (Table 2 and supplementary Table S1).

#### Behaviour: GLM-based analysis for reported number of stimuli – audiovisual weight index

We quantified the influence of the true number of auditory and visual stimuli on the reported (behavioral) auditory or visual numeric estimates using a linear regression model^13^. In this regression model, the reported number of stimuli were predicted by the true number of auditory and visual stimuli separately in the four conditions in the 2 (numeric disparity) × 2 (task relevance) factorial design. The auditory (ß_A_) and visual (ß_V_) parameter estimates quantified the influence of the experimentally defined auditory and visual stimuli on the perceived/decoded number of stimuli for a particular condition. To obtain a relative audiovisual weight index w_AV_, we computed the four-quadrant inverse tangens of the auditory (β_A_) and visual (β_V_) parameters estimates for each of the four conditions (i.e. w_AV_ = atan(β_V_, β_A_)). An audiovisual weight index w_AV_ = 90° indicates purely visual and w_AV_ = 0° purely auditory influence on the reported/decoded number of stimuli.

We performed the statistics on the behavioral audiovisual weight indices using a two (auditory vs. visual report) × two (large vs. small numeric disparity) factorial design based on a likelihood ratio test statistic (LRTS) for circular measures^62^. Similar to an analysis of variance for linear data, LRTS computes the difference in log likelihood functions for the full model that allows differences in the mean locations of circular measures between conditions (i.e, main and interaction effects) and the reduced null model that does not model any mean differences between conditions. To refrain from making any parametric assumptions, we evaluated the effects of task-relevance, numeric disparity and their interaction in the factorial design using randomization tests (5000 randomizations)^63^. To account for the within-subject repeated-measures design at the second random-effects level, randomizations were performed within each participant. For the main effects of numeric disparity and task-relevance, w_AV_ values were randomized within the levels of the non-tested factor^64^. For tests of the numeric disparity x task relevance interactions, we randomized the simple main effects (i.e. (A1B1, A2B2) and (A1B2, A2,B1)) which are exchangeable under the null-hypothesis of no interaction^65^. To test deviations of w_AV_ from specific test angles (e.g. w_AV_ < 90°), we used one-sided one-sample randomization tests in which we flipped the sign of the individual circular distance of w_AV_ from the test angle^66^ and used the mean circular distance as test statistic.

Unless otherwise stated, results are reported at p < 0.05. For plotting circular means of w_AV_ (Fig. 1C and 5C for behavioral w_AV_, Fig. 3C for neural w_AV,_ see multivariate EEG analysis), we computed the means’ bootstrapped confidence intervals (1000 bootstraps).

#### Behaviour: Full-segregation, forced-fusion and Bayesian Causal Inference models

Next, we fitted the full-segregation, the forced-fusion and the Bayesian Causal Inference model with model averaging, model selection and probability matching as decision functions to observers’ behavioural reports. Using Bayesian model comparison, we then assessed which of these models is the best explanation for observers’ reported numeric estimates.

In the following, we will first describe the Bayesian Causal Inference model from which we will then derive the forced-fusion and full-segregation model as special cases. Details can be found in Kording et al. (2007)^1^.

Briefly, the generative model (Fig. 2C) assumes that common (C = 1) or independent (C = 2) causes are determined by sampling from a binomial distribution with the causal prior P(C = 1) = p_common_ (i.e. a priori binding tendency^25^). For a common cause, the “true” number of audiovisual stimuli N_AV_ is drawn from the numeric prior distribution N(μ_P_, σ_P_). For two independent causes, the “true” auditory (N_A_) and visual (N_V_) numbers of stimuli are drawn independently from this numeric prior distribution. We introduced sensory noise by drawing x_A_ and x_V_ independently from normal distributions centered on the true auditory (respectively visual) number of stimuli with parameters σ_A_ (respectively σ_V_). Thus, the generative model included the following free parameters: the causal prior p_common_, the numeric prior’s mean μ_P_ and standard deviation σ_P_, the auditory standard deviation σ_A_, and the visual standard deviation σ_V_. The posterior probability of the underlying causal structure can be inferred by combining the causal prior with the sensory evidence according to Bayes rule:

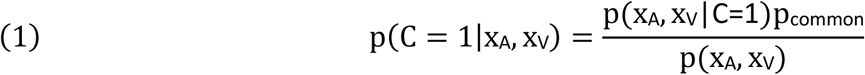

The causal prior quantifies observers’ belief or tendency to assume a common cause and integrate stimuli prior to stimulus presentation. After stimulus presentation, the disparity between the number of beeps and flashes informs the observers’ causal inference via the likelihood term (cf. Fig. 2C). In the case of a common cause (C = 1), the optimal audiovisual numeric estimate 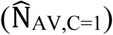 is obtained under the assumption of a squared loss function, by combining the auditory and visual numeric estimates as well as the numeric prior (with a Gaussian distribution of N(μ_P_, σ_P_)) weighted by their relative reliabilities:

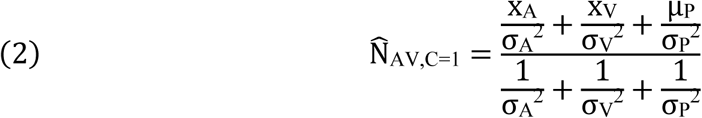

In the case of independent causes (C = 2), the optimal numeric estimates of the unisensory auditory 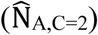 and visual (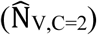) stimuli are independent:

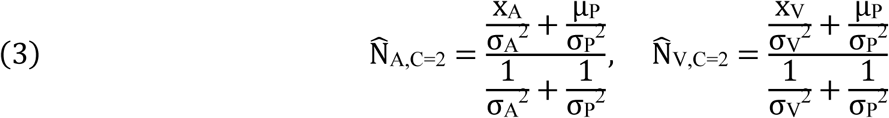

To provide a final estimate of the number of auditory or visual stimuli, the observer is thought to combine the estimates under the two causal structures using various decision functions such as “model averaging,” “model selection,” or “probability matching”^20^. According to “model averaging”, the brain combines the two auditory numeric estimates weighted in proportion to the posterior probabilities of their underlying causal structures:

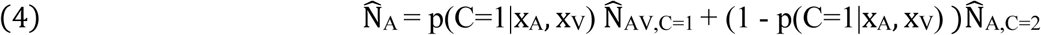

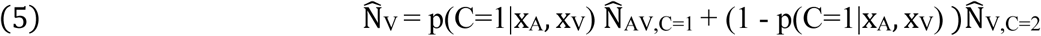

According to the ‘model selection’ strategy, the brain reports the numeric estimate selectively from the more likely causal structure (eq. 6 only for 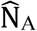):

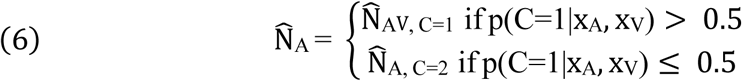

According to ‘probability matching’, the brain reports the numeric estimate of one causal structure stochastically selected in proportion to its posterior probability (eq. 7 only for 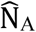):

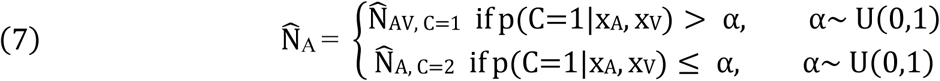

Thus, Bayesian Causal Inference formally requires three numeric estimates 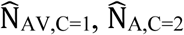, 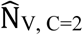 which are combined into a final estimate (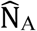 or 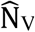, depending on which sensory modality is task-relevant) according to one of the three decision functions.

We evaluated whether and how participants integrate auditory and visual stimuli based on their auditory and visual numeric reports by comparing (i) the full-segregation model that estimates stimulus number independently for vision and audition (i.e. formally, the BCI model with a fixed p_common_ = 0), (ii) the forced-fusion model that integrates auditory and visual stimuli in a mandatory fashion (i.e. formally, the BCI model with a fixed p_common_ = 1) and (iii) the BCI model (i.e. model averaging; Table 2) Because the decisional strategy of ‘model averaging’ outperformed the other decision functions (eq. (4)-(7))) based on Bayesian model comparison at the group level (Supplementary Table. S1), the main report and analysis of the neural data focuses on model averaging.

To arbitrate between the full-segregation, forced-fusion and BCI models, we fitted each model to participants’ numeric reports (Table 2) based on the predicted distributions of the auditory (i.e. the marginal distributions: 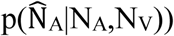 and visual (i.e. 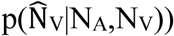) numeric estimates that were obtained by marginalizing over the internal variables x_A_ and x_V_ that are not accessible to the experimenter (for further details of the fitting procedure see Kording et al. (2007)^1^). These distributions were generated by simulating x_A_ and x_V_ 5000 times (i.e. continuous variables sampled from Gaussian distributions) for each of the 32 conditions and inferring 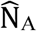 and 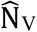 from equations (1)-(5). To link the continuous distributions 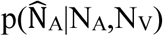 and 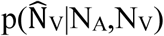 to participants’ categorical auditory or visual numeric reports (i.e. from {1,2,3,4}), we assumed that participants selected the button that is closest to 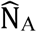 or 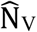 and binned 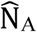 and 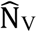 accordingly into a four-bin histogram. From these predicted multinomial distributions (i.e. one for each of the 32 conditions; auditory and visual numeric reports were linked to 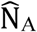 and 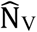, respectively), we computed the log likelihood of participants’ numeric reports and summed the log likelihoods across conditions. To obtain maximum likelihood estimates for the five parameters of the models (p_common_, µ_P_, σ_P_, σ_A_, σ_V_; formally, the forced-fusion and full-segregation models assume p_common_ = 1 or = 0, respectively), we used a non-linear simplex optimization algorithm as implemented in Matlab’s fminsearch function (Matlab R2015b). This optimization algorithm was initialized with 20 different parameter settings that were defined based on a prior grid-search.

We report the results (across-participants’ mean and standard error) of the parameter setting with the highest log likelihood across these initializations (Table 2 and supplementary Table S1). This fitting procedure was applied individually to each participant’s data set for the Bayesian Causal Inference (with three different decision functions), the forced-fusion and the full-segregation models. The model fit was assessed by Nagelkerke’s coefficient of determination^67^ using a null model of random guesses of stimulus number 1-4 with equal probability 0.25. To identify the optimal model for explaining participants’ data, we compared the candidate models using the Bayesian Information Criterion (BIC) as an approximation to the model evidence^68^. The BIC depends on both model complexity and model fit. We performed Bayesian model comparison^69^ at the random effects group level as implemented in SPM12^70^ to obtain the protected exceedance probability (the probability that a given model is more likely than any other model, beyond differences due to chance^71^) for the candidate models.

To generate predictions for the audiovisual weight index based on the Bayesian Causal Inference model (with model averaging), we simulated new x_A_ and x_V_ for 10000 trials for each of the 32 conditions using the fitted BCI model parameters of each participant. For each simulated trial, we computed the BCI model’s i. unisensory visual 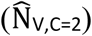, ii. unisensory auditory 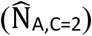 estimates, iii. forced-fusion 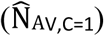, iv. final BCI audiovisual numeric estimate (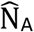 or 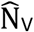 depending on whether the auditory or visual modality was task-relevant) and posterior probability estimate of each causal structure (p(C = 1|x_A_, x_V_)). Next, we used the mode of the resulting (kernel-density estimated) distributions for each condition and participant to compute the model predictions for the audiovisual weight index w_AV_ (Fig. 1C, 5C) and the RDMs (see multivariate EEG analysis, Fig. 2C).

#### EEG data acquisition and preprocessing

EEG signals were recorded from 64 active electrodes positioned in an extended 10-20 montage using electrode caps (actiCap, Brain Products, Gilching, Germany) and two 32 channel DC amplifiers (BrainAmp, Brain Products). Electrodes were referenced to FCz using AFz as ground during recording. Signals were digitized at 1000 Hz with a high-pass filter of 0.1 Hz. Electrode impedances were kept below 25 kOhm.

Preprocessing of EEG data was performed using Brainstorm 3.4^72^ running on Matlab R2015b. EEG data were band-pass filtered (0.25-45 Hz for the main EEG analyses). Eye blinks were automatically detected using data from the FP1 electrode (i.e. a blink was detected if the band-pass (1.5-15 Hz) filtered EEG signal exceeded two times the STD; the minimum duration between two consecutive blinks was 800 ms). Signal-space projectors (SSPs) were created from band-pass filtered (1.5-15 Hz) 400 ms segments centered on detected blinks. The first spatial component of the SSPs was then used to correct blink artifacts in continuous EEG data. Further, all data were visually inspected for artifacts from blinks (i.e. residual blink artifacts after correction using SSPs), saccades, motion, electrode drifts or jumps and contaminated segments were discarded from further analysis (on average 6.4 % ± 0.9 % SEM of all trials discarded). Finally, EEG data were re-referenced to the average of left and right mastoid electrodes and downsampled to 200 Hz. For analysis of event-related potentials (ERPs) and decoding analyses (see below), all EEG data were normalized with a 200 ms prestimulus baseline and were analysed from 100 ms before stimulus onset up to 750 ms after stimulus onset, when the response screen was presented.

#### Preprocessing for multivariate EEG analyses – EEG activity pattern

Single-trial EEG data from the 64 electrodes were binned in time windows of 20 ms. Hence, given a sampling rate of 200 Hz, each 20 ms time window included four temporal sampling points. 64-electrode EEG activity vectors (for each time sample) were concatenated across the four sampling points within each bin resulting in a spatio-temporal EEG activity pattern of 256 features. EEG activity patterns were z scored to control for mean differences between conditions. The first sampling point in the 20 ms time window was taken as the window’s time point in all analyses.

### Overview of EEG analysis

We characterized the neural processes underlying multisensory integration by combining several analysis approaches:

1. Univariate EEG analysis: We identified multisensory integration by testing for audiovisual interactions focusing on the classical ‘sound-induced flash illusion’ conditions, where one flash is presented together with two beeps.
2. Multivariate EEG analysis and neural audiovisual weight index w_AV_: We computed the audiovisual weight index w_AV_ which quantifies the relative influence of the true number of auditory and the visual stimuli on the ‘internal’ numeric estimates decoded from EEG activity patterns using support vector regression (see behavioural analysis above).
3. Multivariate EEG analysis and Bayesian Causal Inference model: We assessed how the numeric estimates obtained from the BCI model, i.e. the unisensory auditory and visual full-segregation, the forced-fusion and the Bayesian Causal Inference estimates (i.e. based on model averaging) are dynamically encoded in EEG activity pattern across post-stimulus time using representation dissimilarity analyses^24^. In supplementary analyses, we also directly decoded the numeric estimates from EEG activity patterns using support vector regression or canonical correlation analyses (Supplementary methods and Fig. S6).
4. Pre-stimulus EEG activity and parameters of the Bayesian Causal Inference model: We investigated whether the power or phase of brain oscillations as measured by EEG before the stimulus onset correlates with the causal prior or the visual precision parameters of the Bayesian Causal Inference model (selectively refitted to trials binned according to their oscillatory power or phase).

#### EEG: Univariate analysis of audiovisual interactions

To assess basic sensory components in ERPs and early audiovisual interactions, we averaged trial-wise EEG data time-logged to stimulus onset into ERPs for audiovisual congruent conditions. We then averaged the ERPs across parietal electrodes (i.e. Cz, CP1, CPz, CP2, P1, Pz, P2; Fig. 1D) or occipital electrodes (i.e. O1, O2, Oz, PO3, POz, PO4; Fig. 1E). To analyze early audiovisual interactions as reported for the sound-induced flash illusion, we computed the difference between audiovisual and the corresponding unisensory conditions (i.e. A_1_V_1_A_2_ - (A_1_A_2_ + V_1_))^21,22^. However, the auditory and visual trials were acquired in separate unisensory runs and may therefore differ in attentional and cognitive context. Further, our experimental design did not include null trials to account for anticipatory effects around stimulus onset and ensure a balanced audiovisual interaction contrast^73^. Hence, these audiovisual interactions need to be interpreted with caution. To test whether the difference wave deviated from zero at the group level, we used a non-parametric randomization test (5000 randomizations) in which we flipped the sign of the individual difference waves and computed a two-sided one-sample t tests as a test statistic^74^. To correct for multiple comparisons across the sampling points, we used a cluster-based correction^75^ with the sum of the t values across a cluster as cluster-level statistic and an auxiliary cluster defining threshold of t = 2 for each time point.

#### EEG: Multivariate GLM-based analysis, decoding accuracy and audiovisual weight index

For each 20 ms time window, we trained linear support-vector regression (SVR) models (libSVM 3.20^76^) to learn the mapping from spatio-temporal EEG activity patterns to the number of flash-beep stimuli of the audiovisually *congruent* conditions (including conditions of auditory and visual report) from all but one run. The SVRs’ parameters (C and ?) were optimized using a grid search within each cross-validation fold (i.e. nested cross-validation). Before training the SVR models, we recoded the stimulus numbers as labels to the range of [−1,1] (i.e. −1 = 1 stimuli; −0.33 = 2 stimuli; 0.33 = 3 stimuli; 1 = 4 stimuli).

This learnt mapping from EEG activity patterns to external number of stimuli was then used to decode the number of stimuli from spatio-temporal EEG activity patterns of the audiovisual *congruent* and *incongruent* audiovisual conditions of the remaining run. In a leave-one-run-out cross-validation scheme, the training-test procedure was repeated for all runs. To account for SNR differences across runs, predicted stimulus numbers were z-scored within each run. The decoded stimulus numbers for the congruent and incongruent conditions were used to assess i. decoding accuracy based on congruent trials only and ii. to compute the audiovisual weight index w_AV_ in subsequent GLM-based analysis approaches (see below).

First, we computed decoding accuracy based selectively on the audiovisual congruent conditions. We decoded stimulus numbers 1-4 at all time points even though the distinctions between high flash and/or beep numbers (e.g. three vs. four) was only possible at later time points. Hence, as expected the decoder was able to discriminate between higher stimulus numbers (e.g. three vs. four stimuli) only after about 250 ms (Fig. 3A). Next, we evaluated the decoder’s accuracy in terms of the Pearson correlation between true and decoded stimulus number selectively in audiovisual congruent conditions (Fig. 3B). We tested whether individual Fisher’s z-transformed correlation coefficients were larger than zero at the group level using a one-sided non-parametric randomization test (sign flip of correlation coefficient in 5000 randomizations) and a cluster-based correction for multiple comparisons across time intervals (as applied to difference waves, see above; cluster-level statistic: sum of the t values in a cluster; auxiliary cluster defining threshold t = 2).

Second, we quantified the influence of the true number of auditory and visual stimuli on the decoded (neural) auditory or visual numeric estimates in a GLM-based analysis approach that was equivalent to our behavioral analysis. In a linear regression model^13^, the decoded number of stimuli was predicted by the true number of auditory and visual stimuli separately for the four conditions in the 2 (numeric disparity) × 2 (task relevance) factorial design (see behavioral analysis for further details). Statistical analysis was also equivalent to the behavioral analysis with the exception that we accounted for multiple comparisons across time using a cluster-based correction (cluster-level statistic: sum of the LRTS values in a cluster; auxiliary cluster defining threshold LRTS = 2). Unless otherwise stated, results are reported at p < 0.05 corrected for multiple comparisons in EEG. For plotting circular means of w_AV_ (Fig. 3C), we computed the means’ bootstrapped confidence intervals (1000 bootstraps).

#### EEG: Multivariate Bayesian Causal Inference and Representational Dissimilarity Matrices

To characterize the neural dynamics of Bayesian Causal Inference, we next investigated whether and when the four numeric estimates of the Bayesian Causal Inference model are represented in EEG activity patterns using support vector regression (i.e. similar to a previous fMRI study^12^), canonical correlation analysis and representational similarity analysis (RSA)^24^. Because these three analysis approaches yield comparable results, we focus in the main manuscript on the RSA (see supplementary materials for the canonical correlation and support vector regression analysis and results, Fig. S6).

To define the representational dissimilarity matrices (RDMs) for the RSA^24^, we computed the pairwise absolute distance between the BCI model’s four numeric estimates, i.e. i. unisensory visual (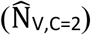 ii. unisensory auditory 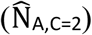 estimates, iii. forced-fusion 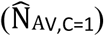, iv final BCI audiovisual numeric estimate (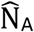 or 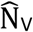 depending on whether the auditory or visual modality were task-relevant) as well as the posterior causal probability across all 32 conditions individually for each participant and then averaged those across participants (Fig. 2C). Likewise, we generated RDMs for the behavioral numeric reports by computing the pairwise absolute distance between the mean numeric reports across all 32 conditions for each participant and then averaged the individual RDMs across participants.

To resolve the evolution of the full-segregation auditory, full-segregation visual, forced-fusion and the BCI estimates in time, we correlated their RDMs with the EEG RDMs. The EEG RDMs were computed as the Mahalanobis distance between single trial spatiotemporal EEG activity patterns for 20 ms time windows over conditions (c.f. decoding analysis above)^77^. More specifically, we computed the Mahalanobis distance from the activity patterns’ variance-covariance matrix using the pattern component modeling toolbox^78^. We quantified the similarity of the RDMs of the numeric estimates of the BCI model (Fig. 2C) with the EEG RDM at each 20 ms time interval using the Spearman’s rank correlation r (Fig. 4A; i.e. correlation of the RDMs’ the upper triangular part). The Fisher’s z-transformed correlation coefficients were tested against zero using a one-sided randomization test (sign flip of correlation coefficient in 5000 randomizations) and a cluster-based correction for multiple comparisons across time intervals (as applied to decoding accuracy, see above).

From the explained variance of the RDMs’ correlation (i.e. r^2^), we computed the Bayesian Information Criterion as an approximation to the model evidence for each estimate and time point^68^ (BIC = n * log(1-r^2^) + 1 * log(n); n = number of EEG activity patterns). We entered these participant-specific model evidences in a random-effects group analysis to compute the protected exceedance probability (SPM12) that one numeric estimate was more likely encoded than any of the other estimates separately for each time interval (Fig. 4B; see above).

#### Time-frequency analysis of the effect of prestimulus oscillations on the causal prior and the visual precision parameters in the Bayesian Causal Inference model

We investigated whether prestimulus oscillatory power or phase over occipital electrodes (i.e. O1, O2, Oz, PO3, POz, PO4)^16,17,26^ is related to the brain’s prior binding tendency as quantified by the causal prior (i.e. p_common_) or the precision of the visual representation (i.e. 1/σ^2^) as estimated in the Bayesian Causal Inference model. We band-pass filtered the continuous EEG data to 0.25-100 Hz with a notch filter at 50 Hz and re-epoched it into trials of −1.5 to 1 s. Using complex Morlet wavelets (as implemented in Brainstorm^72^), we extracted the spectral power and phase of single-trial EEG data from −0.5 s to +0.1s from 6 to 80 Hz in 2 Hz steps with the cycles increasing linearly from 5 to 13 cycles across frequencies^79^. We downsampled the time-frequency representation to 50 Hz (i.e. 38 frequencies × 25 time points). First, based on previous research pointing towards a role of alpha, beta and gamma oscillations in the sound-induced illusion^16,17,26^, we investigated whether the oscillatory power in these bands prior to stimulus onset was correlated with p_common_ or σ_V_. For each point in time-frequency space, we sorted and binned the trials according to their oscillatory power (or phase for alpha frequency) into 10 deciles separately for each of the 32 conditions to control for any condition-specific effects (cf. Fig. 1B)^80,81^. Using maximum likelihood estimation, we refitted selectively the p_common_ (resp. σ_V_) parameter of the BCI model to the multinomial distribution of observers’ numeric reports over the 32 conditions separately for each decile (i.e. based on ∼120 trials), while fixing the remaining four BCI parameters to the parameters obtained from the estimation based on the complete data set (i.e. pooled over the 10 deciles). For each point in time-frequency space, we then computed the correlation between the oscillatory power (averaged across trials within a decile) and the BCI parameter (i.e. p_common_, σ_V_) over the 10 deciles for each participant. At the group level, we tested whether the Fisher’s z-transformed correlation coefficients were significantly different from zero using a randomization test (i.e. sign flip of correlation coefficients in 5000 randomizations; test statistic: two-sided t-tests) and a cluster-based correction for multiple comparisons^75^ separately for the alpha (8-12 Hz), beta (14-28 Hz) and gamma (30-80 Hz) bands (Fig. 5A; cluster-level statistic: sum of the t values in a cluster; auxiliary cluster threshold t = 2). Note that initially, prior to the sort-and-bin approach, all five BCI parameters were re-fitted to the whole data set with valid EEG data. This additional refit was required because additional trials (i.e. 2.9 ± 0.4 % (mean ± SEM) of trials) were rejected due to EEG artefacts in the longer epochs from −1.5 s to 1 s. For illustrational purposes, we also computed the relative weight index w_AV_ for each decile both for the BCI model’s predictions and the participants’ numeric reports averaged across the significant clusters (Fig. 5 C).

Second, based on previous research implicating prestimulus alpha phase in temporal binding^26,27^, we investigated whether the circular mean phase (i.e. averaged across trials within a decile) of alpha oscillations (8-12 Hz) was correlated with the BCI parameters (i.e. p_common_ or σ_V_) over alpha phase deciles using linear-circular correlation^66^. To enable an unbiased group level statistic, we first randomized the assignment between mean circular phase and BCI parameters across the deciles (5000 randomizations) within each participant. Next, we computed the percentile of a participant’s true circular-linear correlation in relation to this participant’s null-distribution of circular-linear correlations. At the group level, we then tested whether the across-participants’ mean percentile was significantly greater than 50% (i.e. the mean percentile under the null hypothesis) using a randomization test (i.e. sign flip of deviation of percentile from 50% in 5000 randomizations; test statistic: one-sided t-tests) and cluster-based correction for multiple comparisons (Fig. 6A; cluster-level statistic: sum of the t values in a cluster; cluster threshold t = 2).

To characterize the modulation of p_common_ by alpha phase, we first fitted a sine and cosine to 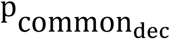 over alpha phase deciles Φdec at 10 Hz individually to each participant’s data, separately for each time point (eq. 8). Thus, we computed the average phase Φdec at 10 Hz across trials for each decile at a particular time point. We then used this average phase Φdec in each decile at 10 Hz to predict 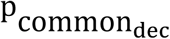 for this particular time point over deciles based on a sinusoidal model:

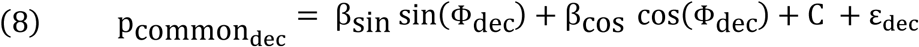

with 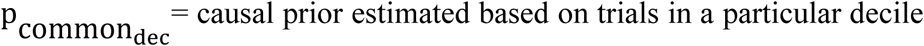; Φdec = across trials average phase in a particular decile; C = constant Crucially, this regression model (i.e. eq. 8) is estimated independently for each time point resulting in βsin and βcos separately for each time point. Thus, eq. 8 characterizes the relationship between alpha phase and pcommon over deciles for a particular time point, so that the phase of this modulation can in principle vary across time (Fig. 6D).

Second, we fitted a more constrained regression model with one single sine and cosine at F = 10 Hz that uses the Φdec t averaged over trials within a particular decile = dec at a time point = t to predict 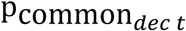. Hence, this model assumes that the modulation of 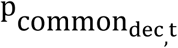 by alpha phase for each time point (i.e. a column in Fig. 6B) evolves slowly over time according to a 10 Hz alpha oscillatory rhythm:

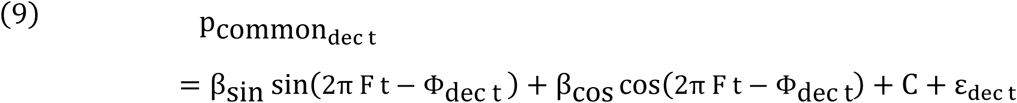

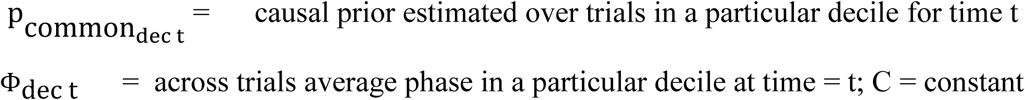

The statistical significance of this model was assessed for the time window of −280 up to −80 ms encompassing the significant cluster (i.e. to include two alpha cycles; Fig. 6A) in each participant with an F test on the residual sum of squares against a reduced model that included only the constant C as a regressor (i.e. df1 = 2; df2 = 107; see Fig. 6B). Next, we assessed whether the phase angle of the alpha oscillation (i.e. Φ_*subject*_ = angle(βcos + i βsin) from equation (9) was consistent across participants and hence deviated significantly from a circular uniform distribution using a Raleigh test^66^. The distribution of phase angles over participants was not significantly different from uniformity, which can be explained by participant-specific cortical folding leading to differences in the orientation of the underlying neural sources. We therefore identified the peak in predicted 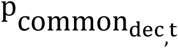 at t = −160 ms in each participant (based on eq. 8), computed the difference in deciles between the participant’s peak decile and the group peak decile (Supplementary Fig. S5B) and then shifted the predicted and observed 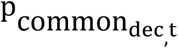 in each participant by this difference across all time points. As a consequence, the adjusted participant’s peak is aligned with the predicted group peak 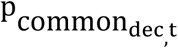 at t = −160 ms; −0.29 π = 52° (Supplementary Fig. S5A, B)^82^. Then we averaged the observed and predicted (cf. eq. 8) pcommon across participants for illustrational purposes (Fig. 6C and D).

#### EEG: The relationship of prior stimulus history, prestimulus alpha power and the causal prior probability

To investigate whether the numeric disparity of prior stimuli influences observers’ causal prior, we sorted current trials according to whether previous trials up to the order of five were of small (≤1) or large (>1) numeric disparity. We selectively refitted the causal prior (holding all other parameters fixed) separately depending on whether the ‘previous trial of a specific order’ was of small or large numeric disparity. We compared the causal prior for small versus large numeric disparity conditions across participants using a 2 (numeric disparity: small vs. large) × 5 (stimulus order: 1, 2, 3, 4, 5 trials back) repeated measures ANOVA. Post-hoc two-sided paired t tests were used to determine up to which trial order previous small numeric disparity led to a larger causal prior as expected.

To investigate whether alpha power (i.e. 8-12 Hz) mediates the effect of prior numeric disparity on the causal prior, we compared alpha power for previous large versus small numeric disparity (i.e. selectively for order one, which had the greatest impact on causal prior) using a randomization test (i.e. sign flip of power difference between previous large vs. small disparity trial in 5000 randomizations; test statistic: two-sided t-tests) and a cluster-based correction for multiple comparisons^75^ (cluster-level statistic: sum of the t values in a cluster; auxiliary cluster threshold t = 2).

Finally, we investigated whether the effect of previous numeric disparity interacts with the correlation between alpha power and the causal prior (i.e. moderation). For this, we first sorted trials according to whether the previous trial (i.e. only order one) was of small or large numeric disparity. We then sorted and binned the trials according to their oscillatory power into 10 deciles separately for previous small versus large prior numeric disparity. We selectively recomputed the causal prior for each decile and assessed the influence of alpha power on causal prior in terms of correlation coefficients separately for previous low and high numeric disparity exactly as in the our initial main analysis on alpha power (see above). Finally, we compared the Fisher’s z-transformed correlation coefficients of alpha power with the causal prior for previous low and high numeric disparity trials in a randomization test (i.e. sign flip of z-transformed correlation differences in 5000 randomizations; test statistic: two-sided t-tests) and cluster-based correction for multiple comparisons (as described above).

#### Assumptions and caveats of multisensory analysis approaches

This study combined several complementary approaches to characterize the neural processes underlying multisensory integration^83^:

Univariate analyses and multisensory interactions: Consistent with previous research^22,23,38^, we identified multisensory integration in terms of audiovisual interactions, i.e. response non-linearities. As discussed in detail in Noppeney (2012)^83^, this approach is limited because single neuron recordings in neurophysiological research have demonstrated that sensory signals are also combined linearly. Linear multisensory integration processes would thus evade interaction analyses. Moreover, interactions computed as AV versus A+V can result if processes are involved for each stimulus component such that the sum of the two unisensory and the multisensory conditions are not matched. For instance, if observers perform a task, decision-and response-preparation-related processes will be counted twice for the sum of the unisensory conditions (i.e. A+V), but be involved only once for the multisensory condition (i.e. AV). Likewise, early putative audiovisual interactions in EEG have been suggested to emerge because of anticipatory ERP effects that precede all stimulus presentations and are therefore counted twice for A+V, but only once for AV (see^73^). Therefore, multisensory interactions should optimally be computed including ‘null events’ to account for non-specific expectation effects (i.e. AV+Null vs. A+V). Further, in our study unisensory and multisensory conditions may differ in attentional context, because auditory and visual conditions were performed in separate experimental runs where either auditory or visual information were task-relevant. Collectively, these factors need to be taken into account when interpreting audiovisual interactions in EEG (or fMRI) responses in our and other studies.

Multivariate decoding: The EEG activity patterns measured across 64 scalp electrodes represent a superposition of activity generated by potentially multiple neural sources located for instance in auditory, visual or higher-order association areas. The extent to which auditory or visual information can be decoded from EEG activity pattern depends therefore inherently not only on how information is neurally encoded by the ‘neural generators’ in source space, but also how these neural activities are expressed and superposed in sensor space (i.e. as measured by scalp electrodes). For example, the number of auditory beeps is perceptually more precisely represented than the number of flashes (based on observers’ behavioral reports, Table 2), suggesting that the brain encodes the timing and number of events with a greater precision in audition than vision. Nevertheless, supplementary decoding analyses in sensor space revealed that the number of unisensory flashes can be more accurately decoded from EEG activity patterns than the number of unisensory beeps (Supplementary Fig. S2). These discrepancies between precision (or accuracy) measured at the behavioral/perceptual level and EEG decoding accuracy at the sensor level may result from differences in neural encoding in source space or how these neural activities are expressed in sensor space (e.g. source orientation, superposition etc.). Potentially, the greater decodability of visual numeric information may contribute to the visual bias we observed for the audiovisual weight index w_AV_ (Fig. 3C) and the dominance of the visual numeric estimates in our decoding analysis based on the estimates of the Bayesian Causal Inference model (Fig. 4A).

In the analysis of the audiovisual weight index w_AV,_ we trained the support vector regression model on the audiovisual congruent conditions pooled over task-relevance to ensure that the decoder was based on activity patterns generated by sources related to auditory, visual and audiovisual integration processes. Moreover, this approach ensures that the effects of task-relevance on the audiovisual weight index w_AV_ cannot be attributed to differences in the decoding model (see^84^ for a related discussion). In a supplementary analysis, we also trained the SVR models separately for visual and auditory report and obtained comparable results (Supplementary Fig. S7) suggesting that our results are immune to this particular choice of decoding model. While the univariate interaction analysis (see above) cannot identify linear response combinations, this multivariate decoding analysis cannot exclude the possibility that auditory and visual stimuli jointly influence EEG activity pattern even though auditory and visual signals are not integrated at the single neuron level.

In our second multivariate analysis approach, we decoded (directly: SVR, canonical correlation analysis; or indirectly: RDM analysis) the numeric estimates of the Bayesian Causal Inference model from EEG activity pattern and then computed the exceedance probability that one numeric estimate was more likely encoded than any other one. The decoding approaches using support vector regression, canonical correlation analysis and representational dissimilarity analysis provided comparable results indicating that our results are robust to the specific decoding approach (Fig. 4 and supplementary Fig. S6). However, given the caveats discussed above (e.g. superposition of EEG activity patterns) and the high correlation between the different numeric estimates in the BCI model, it seems likely that multiple numeric estimates are concurrently represented in the brain even if the exceedance probability is high for only one particular numeric estimate.

## Code availability

The Matlab code to fit the Bayesian Causal Inference model^1^ to the behavioral data is available in the Dryad repository, [PERSISTENT WEB LINK TO DATASETS; UPLOAD WILL FOLLOW UPON ACCEPTANCE OF THE MANUSCRIPT]. Custom Matlab code for the analyses of EEG data are available from the corresponding author on reasonable request.

## Acknowledgements

This study was funded by the University of Tuebingen (Fortüne grant numbers 2292-0-0 and 2454-0-0), the Deutsche Forschungsgemeinschaft (DFG; grant number RO 5587/1-1) and the ERC (ERC-multsens, 309349).

## Author contributions

TR, ACE and UN analyzed the data. TR and UN wrote the manuscript. TR conceived the experiment and collected the data.

## Data availability statement

The datasets generated and analyzed in the current study are available in the Dryad repository, [PERSISTENT WEB LINK TO DATASETS; UPLOAD WILL FOLLOW UPON ACCEPTANCE OF THE MANUSCRIPT].

