## Supplementary Material for "The neural dynamics of hierarchical Bayesian inference in multisensory perception"

### Supplementary Methods

To characterize the neural dynamics of Bayesian Causal Inference, we investigated whether and when the internal estimates of the Bayesian Causal Inference (BCI) model are represented in EEG activity patterns using representational similarity analysis (see main manuscript), support vector regression (SVR) and canonical correlation analysis (CCA). The BCI model includes four internal numeric estimates: the i. unisensory visual ($\widehat{N}_{V,C=2}$), ii. unisensory auditory ($\widehat{N}_{A,C=2}$) estimates, iii. forced-fusion ($\widehat{N}_{AV,C=1}$), iv. final BCI audiovisual numeric estimate ($\widehat{N}_{A}$ or $\widehat{N}_{V}$ depending on whether the auditory or visual modality were task-relevant). Further, we considerd the posterior causal probability across all 32 conditions.

For the CCA approach, we correlated the EEG activity patterns from each 20 ms time window with each of the internal BCI estimates separately, using Matlab's canoncorr function. Because canonical correlations are by definition positive and are not subject to a regularization term (i.e. larger feature spaces lead to larger canonical correlations), we subtracted the mean prestimulus canonical correlation (from -100 to 0 ms) before plotting the first canonical correlation coefficient as a function of time (Supplementary Figure 6A). The Fisher's z-transformed correlation coefficients were tested against zero using a randomization test (sign flip of correlation coefficient in 5000 randomizations) and a cluster-based correction for multiple comparisons across time intervals (as applied for RSA in the main manuscript). From the canonical correlations' Wilks' λ, we computed the Bayesian Information Criterion as an approximation to the model evidence for each numeric estimate and time point ($BIC = n * \log(\lambda) + 1 * \log(n)$; n = number of EEG activity patterns). We entered these participant-specific model evidences into a random-effects group analysis to compute the protected exceedance probability (SPM12) that one numeric estimate was more likely encoded than any of the other estimates separately for each time interval (Supplementary Fig. S6B).

For the SVR approach, we trained SVR models (libSVM 3.20[1]) to learn the mapping from spatio-temporal EEG activity patterns from each 20 ms time window to the internal BCI estimates from all but one run. This learnt mapping was then used to decode the internal BCI estimates from EEG activity pattern of the remaining run. In a leave-one-run-out cross-validation scheme, the training-test procedure was repeated for all runs. The SVRs' parameters (C and ν) were optimized using a grid search within each cross-validation fold (i.e. nested cross-validation). Before training the SVR models, we recoded the internal estimates to the range of [-1,1]. To quantify the decoding accuracy of the internal BCI estimates, we computed the Pearson correlation coefficient between true and decoded internal BCI estimate. The Fisher's z-transformed correlation coefficients were tested against zero (Supplementary Fig. S6C) and the protected exceedance probability for each internal BCI estimates were computed as described for the RSA approach (Supplementary Figure S6D).

### Supplementary Results

*Response time analysis*

To avoid motor-response confounds in EEG signals, participants were instructed to respond only upon the presentation of a response screen, i.e. 750 ms after the onset of the first stimulus (Fig. 1A). As a result, response times (RTs) are less informative, because RTs comprise an unaccountable time window during which participants withhold their response to wait for the response screen. Yet, a 2 (numeric disparity: small ≤ 1 vs. large ≥ 2) x 2 (task relevance: auditory vs. visual report) repeated-measures ANOVA on RTs revealed a significant main effect of numeric disparity ($F_{1, 22} = 14.63$, $p = 0.001$, partial $\eta^2 = 0.399$). In line with the multisensory congruency effect on RTs[2], smaller disparity led to faster RTs (Supplementary Figure S1). Critically, we also observed an interaction between numeric disparity and task relevance ($F_{1, 22} = 5.74$, $p = 0.026$, partial $\eta^2 = 0.207$). Observers were slower to report the number of flashes than beeps particularly for large (relative to small) disparity trials. In other words, observers were slower to report the less precise visual estimate particularly when they had to segregate sensory signals.

**Supplementary Figures**

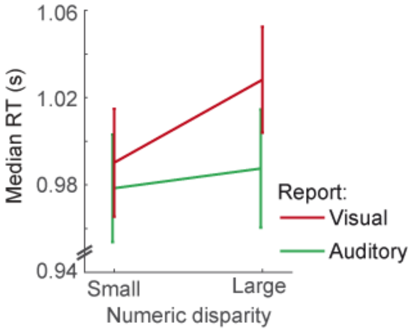


**Supplementary Figure S1. Reaction times.** Reaction times (RT; across-participants mean ± SEM; n = 23) are shown as a function of numeric disparity (small: ≤ 1 vs. large: > 1) and task relevance (auditory vs. visual report). Reaction times were computed as the mean across participants' median reaction times. Note that reaction times have to be interpreted with caution because participants had to withhold their report until the response screen was presented, i.e. 750 ms after the onset of the first stimulus (cf. Fig. 1A).

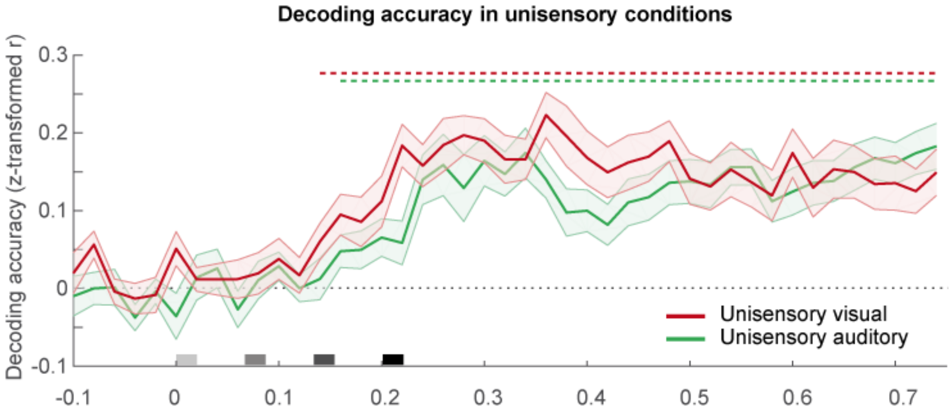


**Supplementary Figure S2. Decoding accuracy in unisensory visual and auditory conditions.** Decoding accuracy (Fisher's z-transformed correlation; across-participants mean ± SEM; n = 23) of the unisensory decoders as a function of time. Auditory (resp. visual) decoders were trained to decode the stimulus number from EEG activity patterns from unisensory auditory (resp. visual) stimulation. Decoding accuracy was computed as Pearson correlation coefficient between true and decoded stimulus number in auditory (resp. visual) conditions. The horizontal dashed lines indicate significant ($p < 0.05$) clusters of decoding accuracy (one-sided cluster-based corrected randomization $t_{22}$ test). Stimulus onsets are shown on the x axis. red = unisensory visual; green = unisensory auditory.

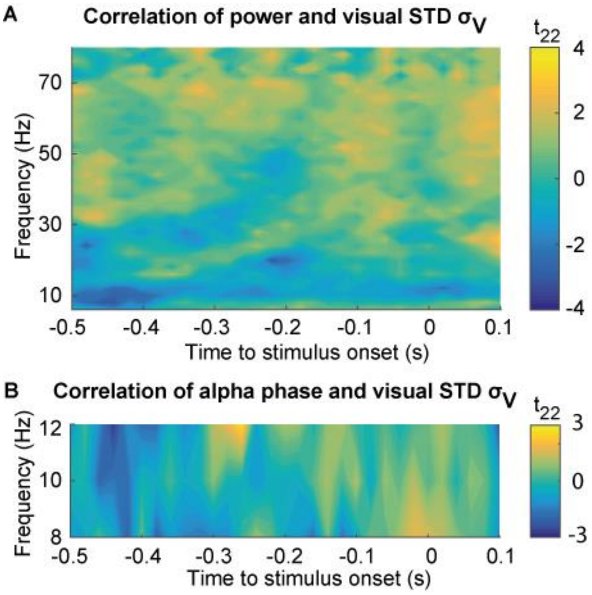


**Supplementary Figure S3. Effect of prestimulus oscillations on the BCI model's visual standard deviation ($\sigma_V$) in occipital electrodes. (A)** Time-frequency t-value map (n = 23) for the correlation between $\sigma_V$ (A) and the oscillatory power deciles averaged across occipital electrodes. No significant cluster (p>0.05; two-sided cluster-based corrected randomization $t_{22}$ test) was found**. (B)** Time-frequency t-value map for the circular-linear correlation between $\sigma_V$ and the phase deciles of alpha oscillations averaged across occipital electrodes. No significant cluster (p >0.05) was found.

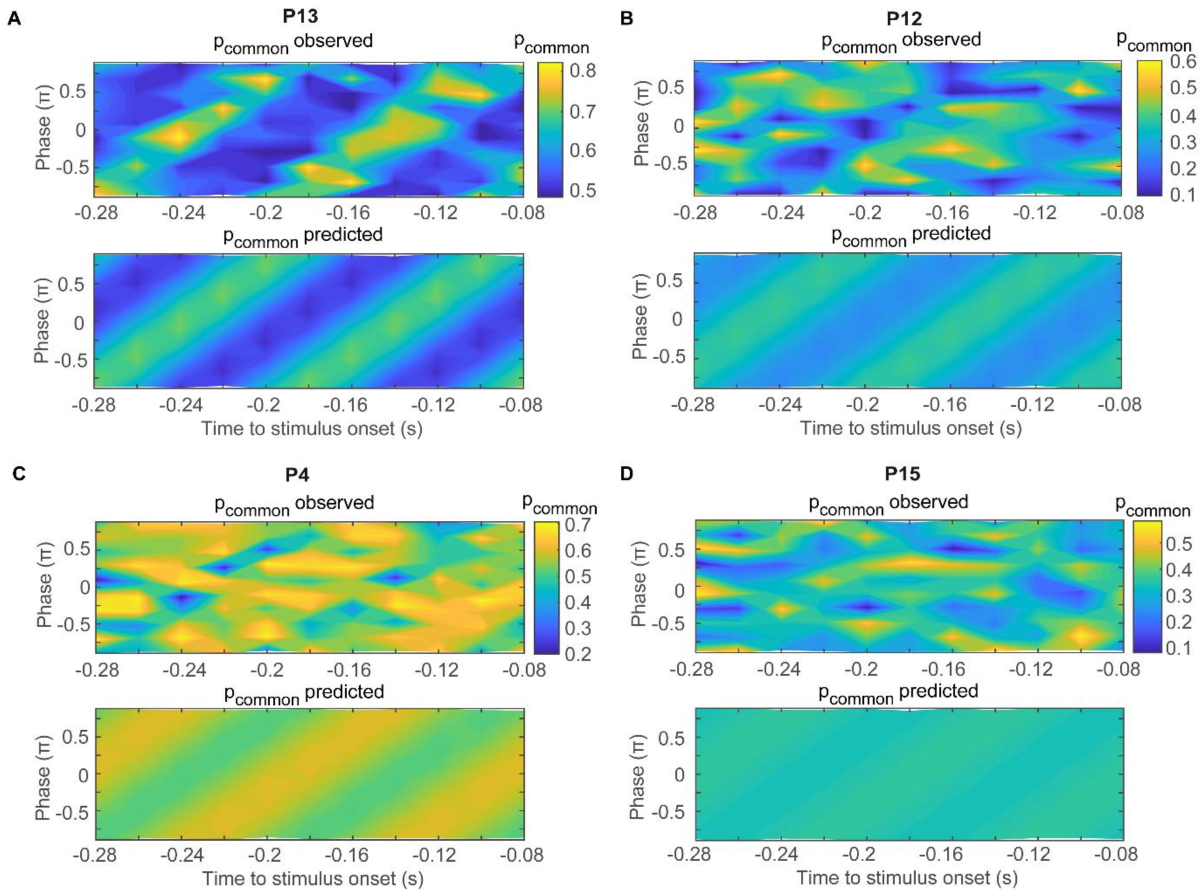


**Supplementary Figure S4. Effect of prestimulus alpha phase on the BCI model's causal prior ($p_{common}$) in occipital electrodes in four representative participants. (A)** Upper panel: The observed $p_{common}$ is shown coded in color for the time window from -280 up to -80 ms, as a function of 10 Hz alpha phase. Lower panel: The predicted $p_{common}$ is based on the parsimonious sinusoidal model of $p_{common}$ (i.e. eq. 9) which assumes that the modulation of $p_{common}$ by alpha phase evolves at 10 Hz. In this participant, the model fit is highly significant ($F_{2, 107} = 51.0$, $p < 0.001$; F test). **(B)** Data as shown in (A) for a participant with significant, but weaker model fit ($F_{2, 107} = 7.0$, $p = 0.001$). **(C)** Data as shown in (A) for a participant with significant, but weaker model fit ($F_{2, 107} = 4.2$, $p = 0.018$). **(D)** Data as shown in (A) for a participant with a non-significant model fit ($F_{2, 107} = 0.4$, $p = 0.659$).

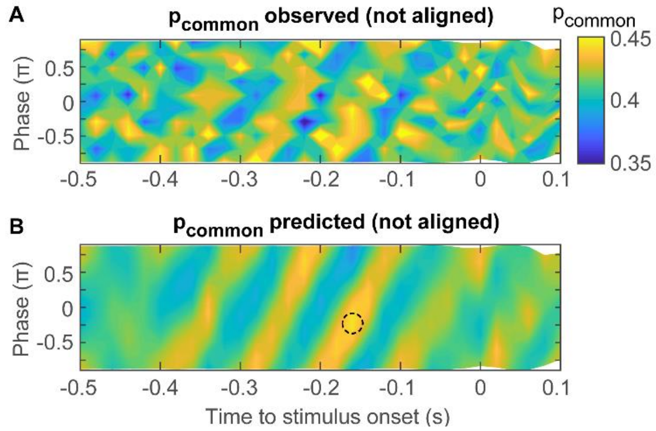


**Supplementary Figure S5. Effect of prestimulus alpha phase on the BCI model's causal prior ($p_{common}$) in occipital electrodes, when data is not phase-aligned.** Across-participants' mean (n = 23) observed **(A)** and predicted **(B)** $p_{common}$ is shown coded in color as a function of time and 10 Hz alpha phase. The phases of the observed and predicted $p_{common}$ were *not* aligned across participants before averaging across participants. The dashed circle in (B) indicates the peak effect to which individual data is phase-aligned in Fig. 6C and D. The predicted $p_{common}$ (B) is based on the sinusoidal models that were fit to $p_{common}$ over alpha phase deciles independently for each time point and participant (eq. 8).

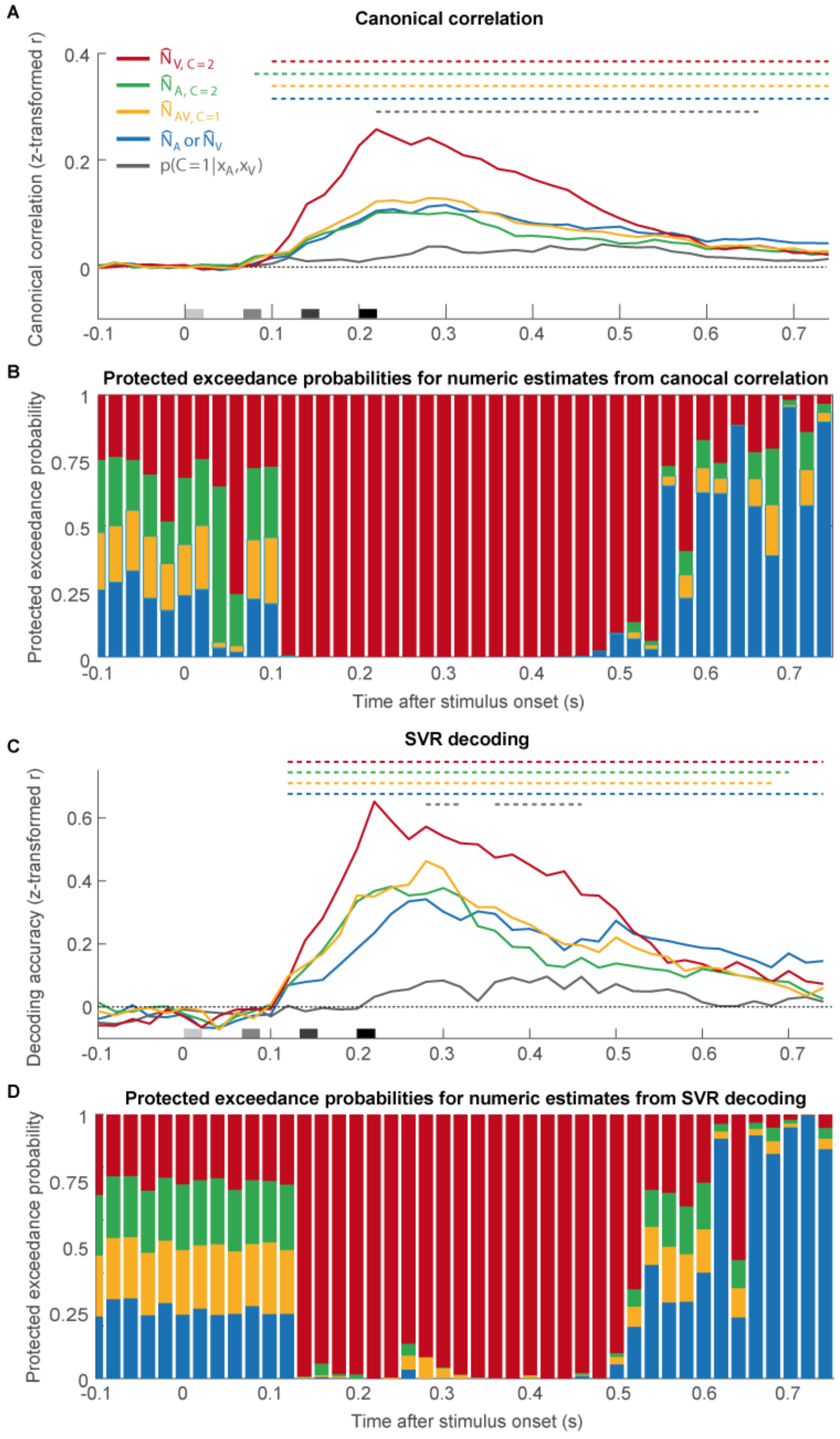


**Supplementary Figure S6. Linking the representational geometry of the BCI models' estimates and EEG patterns by canonical correlation analysis and support-vector regression (SVR) decoding. (A)** Canonical correlation coefficients (across-participants mean of Fisher's z-transformed r; n = 23) as a function of time. The canonical correlation coefficients were computed between EEG activity patterns and separately for each of the BCI model's internal estimates: i. the unisensory visual ($\hat{N}_{V,C=2}$), ii. the unisensory auditory ($\hat{N}_{A,C=2}$) estimates under the assumption of independent causes (C = 2), iii. the forced-fusion estimate ($\hat{N}_{AV,C=1}$) under the assumption of a common cause (C = 1) and iv. the final BCI estimate ($\hat{N}_A$ or $\hat{N}_V$ depending on the sensory modality that is task-relevant) that averages the task-relevant unisensory and the reliability-weighted estimate by the posterior probability estimate of each causal structure ($p(C = 1|x_A, x_V)$). Color-coded horizontal dashed lines indicate clusters of significant correlation coefficients ($p < 0.05$; one-sided cluster-based corrected randomization $t_{22}$ test). Note that the mean of the canonical correlations in the prestimulus time window (i.e. -100 ms to 0 ms) was subtracted from the canonical correlations. Stimulus onsets are shown along the x axis. **(B)** To identify which of the four numeric BCI estimates is most likely represented in the EEG activity patterns, we computed the protected exceedance probability (i.e. the probability that a given variable is more likely encoded in the EEG patterns than any other variable, beyond differences due to chance) for each of the four numeric BCI estimates as a function of time. The length of an estimate's bar indicates the estimate's protected exceedance probability (n.b.: the y axis indicates protected exceedance probabilities cumulated over the BCI estimates). **(C)** Decoding accuracy (Fisher's z-transformed correlation; across-participants mean ± SEM) of the SVR decoders as a function of time. Decoding accuracy was computed as Pearson correlation coefficient between true and the BCI model's internal estimates that were decoded from EEG activity patterns using SVR models trained separately for each numeric estimate. Significant clusters shown as in (A). **(D)** The protected exceedance probability that one numeric estimate, decoded with SVR, is more likely encoded in the EEG pattern than any other of the four numeric BCI estimates as a function of time, shown as in (B).

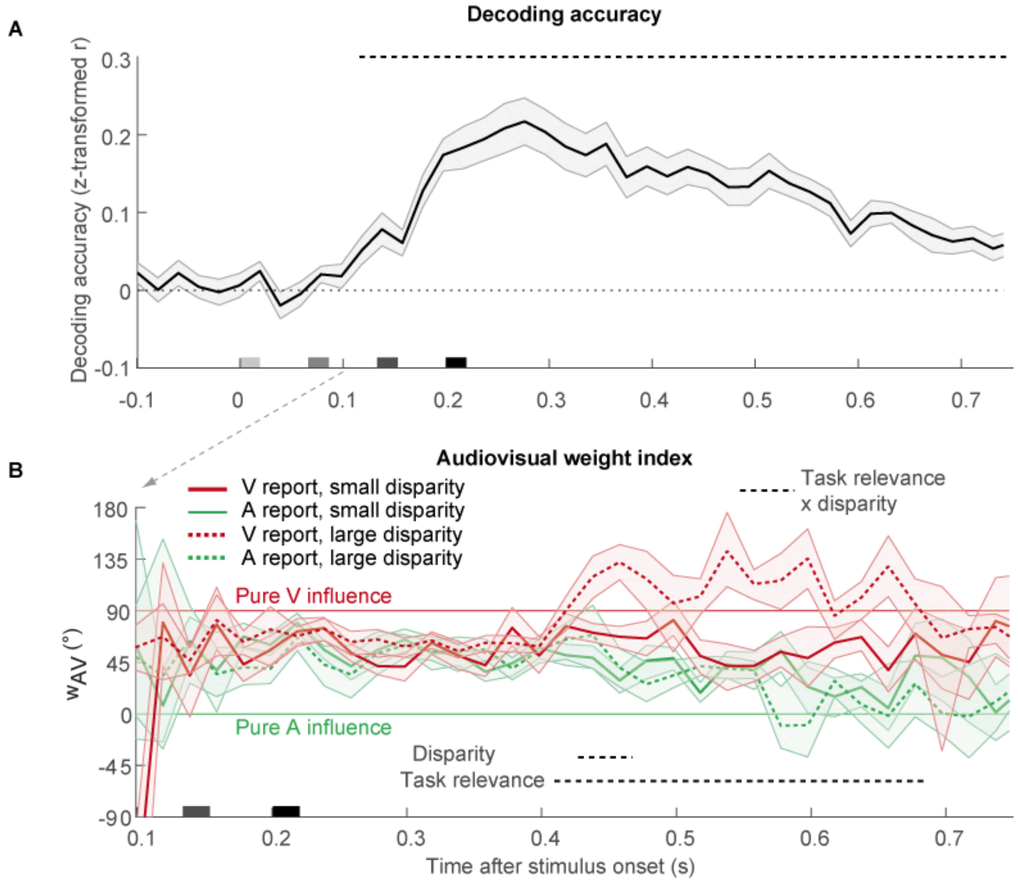


**Supplementary Figure S7. Decoding accuracy and relative audiovisual weight index for decoders trained separately for audiovisual conditions under auditory versus visual report. (A)** Decoding accuracy (Fisher's z-transformed correlation; across-participants mean ± SEM, n = 23) of the decoders as a function of time. Decoding accuracy was computed as Pearson correlation coefficient between true and decoded stimulus number in audiovisual congruent conditions. The horizontal dashed line indicates significant ($p < 0.05$) clusters of decoding accuracy (one-sided cluster-based corrected randomization $t_{22}$ test). Stimulus onsets are shown on the x axis. **(B)** Relative weight index $w_{AV}$ (across-participants circular mean and bootstrapped 68% CI) of the decoders' predicted stimulus number as a function of numeric disparity (small vs. large), task relevance (visual (V) vs. auditory (A) report) and time. $w_{AV} = 90°$ for purely visual and $w_{AV} = 0°$ for purely auditory influence. Horizontal dashed lines indicate significant clusters with effects of numeric disparity, task relevance and their interaction ($p < 0.05$; cluster-based corrected randomization test based on a LRTS statistic).

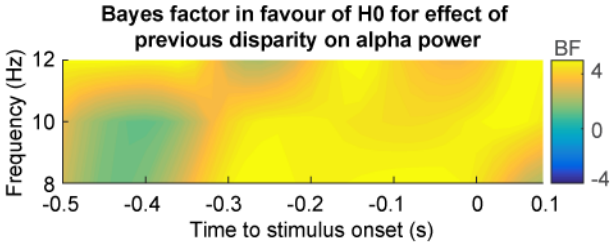


**Figure S8. Bayes factor (BF) in favor of no effect (H0) of the previous disparity on alpha power.** Time-frequency BF-value map (n = 23, averaged over occipital electrodes) shows substantial evidence (i.e., BF > 3.2[3]) in favor of no difference in prestimulus alpha power between small versus large previous numeric disparity (i.e. trial order 1). BFs were computed from the time-frequency t-value map shown in Fig. 7C[4].

### Supplementary table

**Table S1. Results of the Bayesian model comparison of the BCI model with three difference decision functions (model averaging, model selection and probability matching).**

| | $p_{common}$ | $\mu_P$ | $\sigma_P$ | $\sigma_A$ | $\sigma_V$ | $R^2$ | relBIC | pEP | % win |
|---|---|---|---|---|---|---|---|---|---|
| Model averaging | 0.42+0.05 | 2.26+0.20 | 2.34+0.29 | 0.53+0.03 | 1.11+0.23 | 0.875±0.012 | 0.00 | 0.773 | 56.5 |
| Model selection | 0.47+0.05 | 2.35+0.18 | 1.86+0.31 | 0.54+0.03 | 0.82+0.06 | 0.866+0.01.3 | 295.87 | 0.149 | 30.4 |
| Prob. matching | 0.41+0.05 | 2.37+0.17 | 2.06+0.30 | 0.52+0.03 | 1.39+0.46 | 0.869+0.012 | 225.59 | 0.077 | 13.0 |

Note: $p_{common}$, causal prior; $\mu_P$, mean of the numeric prior; $\sigma_P$, standard deviation of the numeric prior; $\sigma_A$, standard deviation of the auditory likelihood; $\sigma_V$, standard deviation of the visual likelihood; $R^2$, coefficient of determination; relBIC, Bayesian information criterion at the group level, i.e. subject-specific BICs summed over all subjects (BIC = LL − 0.5 M ln(N), LL = log likelihood, M = number of parameters, N = number of data points) of a model relative to the Bayesian Causal Inference model with “model averaging” as decision function (n.b. a smaller relBIC indicates that a model provides a better explanation of our data); pEP, protected exceedance probability, i.e. the probability that a given model is more likely than any other model, beyond differences due to chance); % win, percentage of participants in which a model won the within-participant model comparison based on BIC.
